# ARMC9 and TOGARAM1 define a Joubert syndrome-associated protein module that regulates axonemal post-translational modifications and cilium stability

**DOI:** 10.1101/817213

**Authors:** Brooke L. Latour, Julie C. Van De Weghe, Tamara D. S. Rusterholz, Stef J.F. Letteboer, Arianna Gomez, Ranad Shaheen, Matthias Gesemann, Megan E. Grout, Jeroen van Reeuwijk, Sylvia E.C. Van Beersum, Caitlin V. Miller, Jennifer C. Dempsey, Heba Morsy, Michael J. Bamshad, Deborah A. Nickerson, Stephan C.F. Neuhauss, Karsten Boldt, Marius Ueffing, Fowzan S. Alkuraya, Ruxandra Bachmann-Gagescu, Ronald Roepman, Dan Doherty

**Affiliations:** Department of Human Genetics and Radboud Institute for Molecular Life Sciences, Radboud University Medical Center, Geert Grooteplein Zuid 10, 6525 GA Nijmegen, The Netherlands; Department of Pediatrics, University of Washington, Seattle, WA, 98195 USA; Institute of Medical Genetics, University of Zurich, 8952 Schlieren, Switzerland; Institute of Molecular Life Sciences, University of Zurich, 8057 Zürich, Switzerland; Department of Genetics, King Faisal Specialist Hospital and Research Center, Riyadh, Saudi Arabia; University of Washington Center for Mendelian Genomics, Seattle, WA 98195, USA.; Department of Human Genetics, Medical Research Institute, Alexandria University, Alexandria, Egypt; Department of Genome Sciences, University of Washington, Seattle, WA 98195, USA; Medical Proteome Center, Institute for Ophthalmic Research, University of Tuebingen, 72074 Tuebingen, Germany; Department of Anatomy and Cell Biology, College of Medicine, Alfaisal University, Riyadh 11533, Saudi Arabia; Center for Integrative Brain Research, Seattle Children’s Research Institute, Seattle, WA, 98101 USA

## Abstract

Joubert syndrome (JBTS) is a recessive neurodevelopmental ciliopathy, characterized by a pathognomonic hindbrain malformation. All known JBTS-genes encode proteins involved in the structure or function of primary cilia, ubiquitous antenna-like organelles essential for cellular signal transduction. Here, we use the recently identified JBTS-associated protein ARMC9 in tandem-affinity purification and yeast two-hybrid screens to identify a novel ciliary module composed of ARMC9-TOGARAM1-CCDC66-CEP104- CSPP1. *TOGARAM1*-variants cause JBTS and disrupt its interaction with ARMC9. Using a combination of protein interaction analyses and characterization of patient-derived fibroblasts, CRISPR/Cas9-engineered zebrafish and hTERT-RPE1 cells, we demonstrate that dysfunction of *ARMC9* or *TOGARAM1* results in short cilia with decreased axonemal acetylation and glutamylation, but relatively intact transition zone function. Aberrant serum-induced ciliary resorption and cold-induced depolymerization in both *ARMC9* and *TOGARAM1* patient cells lines suggest a role for this new JBTS-associated protein complex in ciliary stability.

## Introduction

Ciliopathies are a heterogeneous class of disorders that arise from defects in the structure or function of the primary cilium (1), a highly specialized microtubule-based sensory organelle that protrudes from the surface of most eukaryotic cell types (2). Joubert syndrome (JBTS) is a recessive, genetically heterogeneous, neurodevelopmental ciliopathy, defined by a distinctive brain malformation, recognizable as the “molar tooth sign” (MTS) (3) in axial magnetic resonance imaging through the midbrain-hindbrain junction. Affected individuals have hypotonia, ataxia, abnormal eye movements, and cognitive impairment. Additional features can occur, such as retinal dystrophy, fibrocystic kidney disease, liver fibrosis, polydactyly, and coloboma. To date, variants in more than 35 genes have been causally linked to JBTS, but the genetic diagnosis has not been identified in all patients, and the disease mechanism remains unclear (4).

All JBTS-associated proteins to date function in and around the primary cilium, but their specific dysfunction can affect a wide variety of cellular processes, including cilium formation, resorption, tubulin post-translational modifications, membrane phosphatidylinositol composition, ciliary signaling pathways, actin cytoskeleton dynamics and DNA damage response signaling (5–10). Many JBTS proteins act together in modules that localize to specific subdomains of the ciliary compartment and disruption in the composition, architecture, or function of these subdomains causes disease (4).

The core of the cilium is composed of nine microtubule doublets forming the ciliary axoneme, which is anchored to the cell by the basal body, a modified centriole. The axonemal microtubules undergo a range of post-translational modifications including polyglutamylation and acetylation which are important for the structure and function of the cilium (11–13). The ciliary membrane has a distinct protein and lipid distribution that differs from the contiguous plasma membrane. This unique composition is achieved in part by the transition zone (TZ) that connects the axoneme to the membrane and acts as a partition. Approximately half of the known JBTS proteins, including RPGRIP1L (JBTS7) (14) and CC2D2A (JBTS9) (15), assemble into multi-protein complexes at the ciliary TZ where they are thought to organize the molecular gate that regulates ciliary protein entry and exit (16); and dysfunction of TZ organization is thought to play a key role in JBTS. Another subset of JBTS-associated proteins, including ARL13B (JBTS8) and INPP5E (JBTS1) (6), associate with the ciliary membrane beyond the TZ. These proteins are thought to play a role in regulation of signaling pathways such as Hh signaling via maintenance of ciliary lipid composition (17, 18). Different JBTS-associated proteins have been found to function at the basal body or distal segment/tip (4). CSPP1 (JBTS 21) (19) and CEP104 (JBTS25) (20) were mainly detected at the centrosomes and ciliary basal bodies, however their exact molecular function, and how defects in these proteins lead to JBTS, are less well understood. CEP104 localizes to the ciliary tip during ciliogenesis, where it is required for structural integrity in the motile cilia of *Chlamydomonas* and *Tetrahymena* (21, 22). Mutations in the gene encoding the ciliary tip kinesin KIF7 (JBTS12) cause JBTS, which were linked to defects in tubulin acetylation and Hedgehog signaling (23).

Recently, we identified mutations in the gene encoding armadillo repeat motif containing 9 (ARMC9) in individuals with JBTS (JBTS30). ARMC9 localizes to centrioles (27) and the proximal portion of the cilium (28) in mammalian cilia. *ARMC9* transcript levels are upregulated with induction of ciliogenesis and *armc9* dysfunction in zebrafish yields typical ciliopathy phenotypes (27). As ARMC9 has not been identified as a component of the ciliary JS-associated protein modules mentioned above, we here use ARMC9 as a bait in protein interaction screens. This delineates a microtubule-associated ciliary protein module containing multiple other JBTS-associated proteins (ARMC9, CEP104, CSPP1, RPGRIP1L, and CEP290) and two ciliary proteins (TOGARAM1 and CCDC66) not previously implicated in JBTS. Strikingly, we identify pathogenic *TOGARAM1* variants in three families affected by JBTS. To decipher the function of this protein module and assess its role in the pathology of JBTS, we map the interaction domains and evaluate cellular defects in cultured human cells and zebrafish mutants. We find that the proteins of this module, previously shown to associate with microtubules (24, 29–31), are required for appropriate post-translational modification of ciliary microtubules and cilium stability, processes that are compromised by loss of both *ARMC9* and *TOGARAM1*.

## Results

### Identification of a novel protein module implicated in JBTS

To define ARMC9-containing protein complexes, we performed protein interaction screens. Using strep - FLAG epitope-tagged ARMC9 expressed in HEK293T cells, followed by tandem affinity purification (TAP) and subsequent mass spectrometry, we identified 116 candidate ARMC9 interactors, including the known JBTS-associated proteins CEP290, CEP104, CSPP1 (Figure 1A, B; Supplemental table 1). Within the data set, ARMC9, CEP104, CCDC66, and TOGARAM1/Crescerin-1 were chosen for further characterization as they are microtubule-associated proteins previously implicated in cilium function (26, 28, 29). Co-immunoprecipitation (co-IP) confirmed the interaction between TOGARAM1 and ARMC9 (Figure 1C). Subsequent TAP experiments using tagged TOGARAM1 and CCDC66 confirmed ARMC9 as their shared binding partner and extended the network to include several other ciliary proteins (Supplemental table 2, Supplemental table 3). For TOGARAM1, these candidate interactors included ciliary proteins ARMC9, CEP104, IFT74, IFT172, PLK1, and PRPF31, (Figure 1A, B; Supplemental table 2) while for CCDC66, they included ARMC9 and DYNLL1 (Figure 1A; Supplemental table 3).

**Figure 1.**
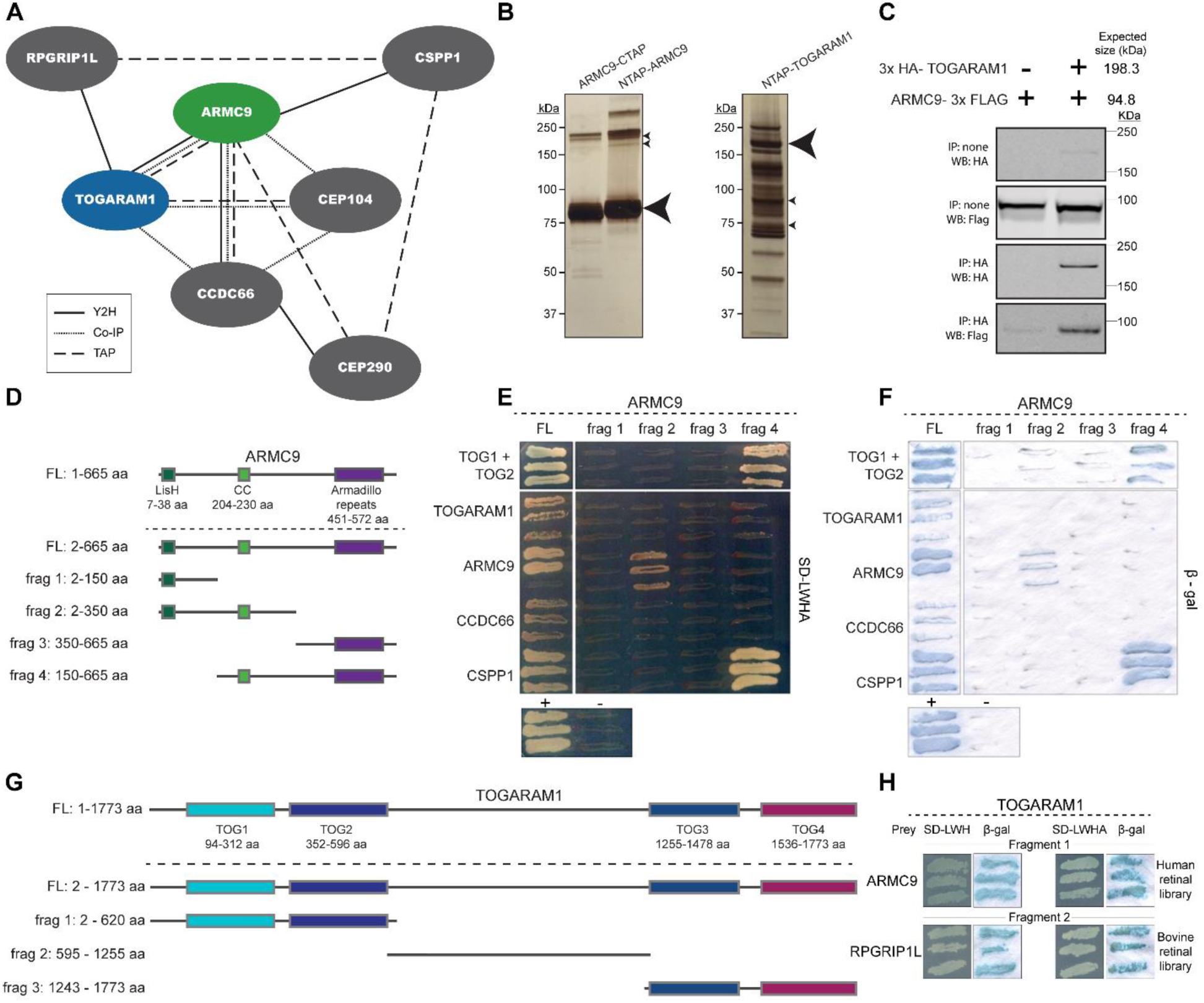
ARMC9 associates with TOGARAM1 in a novel ciliary module. **(A)** ARMC9 interacts with TOGARAM1 as confirmed by tandem affinity purification (TAP), dashed line, and yeast to-hybrid (Y2H) screens, solid line, validation was subsequently performed using co-immunoprecipitation (co-IP), dotted line. **(B)** Silver stain gel of C-terminally and N-terminally SF-TAP tagged ARMC9 (left, large arrow, 80 kDa) and N-terminally SF-TAP tagged TOGARAM1 (right, large arrow, 200 kDa) post protein purification. The small arrows indicate the expected protein bands of two TOGARAM1 isoforms (195.6 kDa and 189.4 kDa) in the ARMC9 TAP purification, and two endogenous ARMC9 isoforms (91.8 kDa and 75.7 kDa) in the TOGARAM1 TAP purification. **(C)** Co-IP of 3xHA-TOGARAM1 and ARMC9-3xFlag. IMCD3 Flp-In cells constitutively expressing ARMC9-3xFlag were transfected with 3xHA-TOGARAM1. Western blot analysis post HA purification indicates the presence of FLAG tagged ARMC9 confirming the interaction. **(D)** Diagram of the ARMC9 full length and fragments used as baits in Y2H bovine and human retinal cDNA library screens. Domains indicated are the predicted Lissencephaly type-1-like homology motif (LisH, 7-38 aa), a coiled-coil domain (cc, 204-230 aa) and the armadillo repeats-containing domain (armadillo, 451- 572 aa). **(E)** The resulting interactors found in the screens are shown in a direct interaction analysis grid using full length prey constructs. Selection of strains co-expressing bait and prey constructs was performed on quadruple knockout SD medium (SD-LWHA). The top row displays yeast colony growth when using fragment 1 of TOGARAM1 containing the TOG1 and TOG2 domains used as prey. (**F)** β- Galactosidase activity assay of **(E)** confirming the interactions. **(G)** Schematic of TOGARAM1 full length and fragments used in a Y2H bovine and human retinal cDNA screen and resulting interactors. **(H)** TOGARAM1 screen results were further validated in a Y2H directed interaction analysis on triple (SD-LWH) and quadruple (SD-LWHA) knockout media.

To identify direct binding partners, we employed full-length and 4 fragments of ARMC9 (Figure 1D) in a GAL4-based yeast two-hybrid (Y2H) interaction trap screen, utilizing two validated prey retinal cDNA libraries that were generated via random or oligo-dT primers (30). Using full-length ARMC9 as a bait, we identified a total of four binary interactors, including ARMC9 itself (suggesting a propensity to multimerize) and the three core module proteins we previously identified by ARMC9 TAP: TOGARAM1, CCDC66, and CSPP1. Validation of these interactions and evaluation of the interacting domains was performed by Y2H co-expression. This assay indicated that the potential self-binding propensity of ARMC9 is mediated by fragment 2 containing the N-terminal 350 amino acid stretch containing the lissencephaly type-1-like homology motif (LisH) and coiled-coil domains, while TOGARAM1 and CSPP1 associate with fragment 4 (150-665 aa) containing the coiled-coil domain and the armadillo repeats domain (Figure 1E, F).

We also used full-length and 3 fragments of TOGARAM1 in parallel screens (Figure. 1G), which confirmed the direct interaction between TOGARAM1 and ARMC9 (Figure 1H), and yielded five additional candidate interactors: 4 without a known role in the cilium (KIFC3, TRAF6, PDE4DIP, and CCDC150) (data not shown), and the JBTS-associated transition zone protein RPGRIP1L (JBTS7) (Figure 1H). The interaction with ARMC9 was mapped to the N-terminal portion of TOGARAM1 (fragment 1) containing the TOG1 and TOG2 domains, while RPGRIP1L bound to the linker region (fragment 2) between the TOG2 and TOG3 domain (Figure 1H).

To further validate the ARMC9 interactors observed in TAP and Y2H experiments, we performed co-IP of ARMC9 with the module components TOGARAM1, CEP104, CCDC66, and CSPP1 (Supplemental Figure 1A). The results confirmed the interaction of ARMC9 with TOGARAM1, CCDC66, and CEP104 but not CSPP1. Additionally, we performed PalmMyr-CFP mislocalization assays to further confirm the interaction of TOGARAM1 and ARMC9. The PalmMyr-CFP assay utilizes a PalmMyr tag which provides sites for palmitoylation and myristoylation. The palmitoylation and myristoylation forces the tagged protein to (mis)localize to the cell membrane (31). Due to the presence of the cyan fluorescent protein (CFP) fusion, this mislocalization can be visualized using fluorescence microscopy. We transfected PalmMyr-CFP tagged ARMC9 and mRFP-tagged TOGARAM1 into human telomerase reverse transcriptase retina pigmented epithelium (hTERT-RPE1) cells, alone and in combination to assess the interaction between ARMC9 and TOGARAM1. Cells transfected with PalmMyr-CFP-ARMC9 alone showed diffuse localization across the cell membrane (Supplemental Figure 1B), while mRFP-TOGARAM1 alone localized along microtubules (Supplemental Figure 1C). Co-expression yielded complete co-localization along microtubules despite the PalmMyr tag, a double membrane anchor, to ARMC9 (Supplemental Figure 1D), thereby indicating a direct interaction of the two proteins and the strong microtubule binding affinity of TOGARAM1.

### *TOGARAM1* variants cause JBTS in humans

Next, we investigated whether our interactome dataset could be used to identify new JBTS-associated genes. We cross-referenced the ARMC9-interactome data with DNA sequence data from our cohort of >600 families affected by JBTS (32). We first evaluated exome sequence data for 53 individuals in 51 families without variants in known JBTS genes. We identified one individual (UW351-3 in Table 1), an affected fetus, with bi-allelic, missense variants in *TOGARAM1* (Figure 2A). These variants (c.1124T>C; p.Leu375Pro and c.3931C>T; p.Arg1311Cys) are rare in gnomAD v2.1 (33) and predicted to be deleterious by combined annotation dependent depletion (CADD) (34) (Table 1). To identify additional families, we used small-molecule molecular inversion probe capture followed by next generation sequencing on 534 additional individuals in the same cohort and identified another individual (UW360-3 in Table 1) with a nonsense variant (c.1084C>T; p.Gln362*) on one allele (Figure 2A) and a possible multi-exon deletion event on the other allele, initially identified as a region of low coverage. We confirmed a 12 kb deletion using a custom comparative genomic hybridization array, and fibroblast cDNA sequencing revealed deletion of exons 4-7, which encode a portion of the linker region between the TOG2 and TOG3 domains (Figure 2B, C). In parallel, exome sequencing in a cohort of 350 families with various ciliopathies identified a homozygous (c.1102C>T; p.Arg368Trp) (Figure 2A) variant in the child of consanguineous parents (13DG1578 in Table 1). This variant has not been reported in population databases and is predicted to be deleterious by CADD.

**Figure 2.**
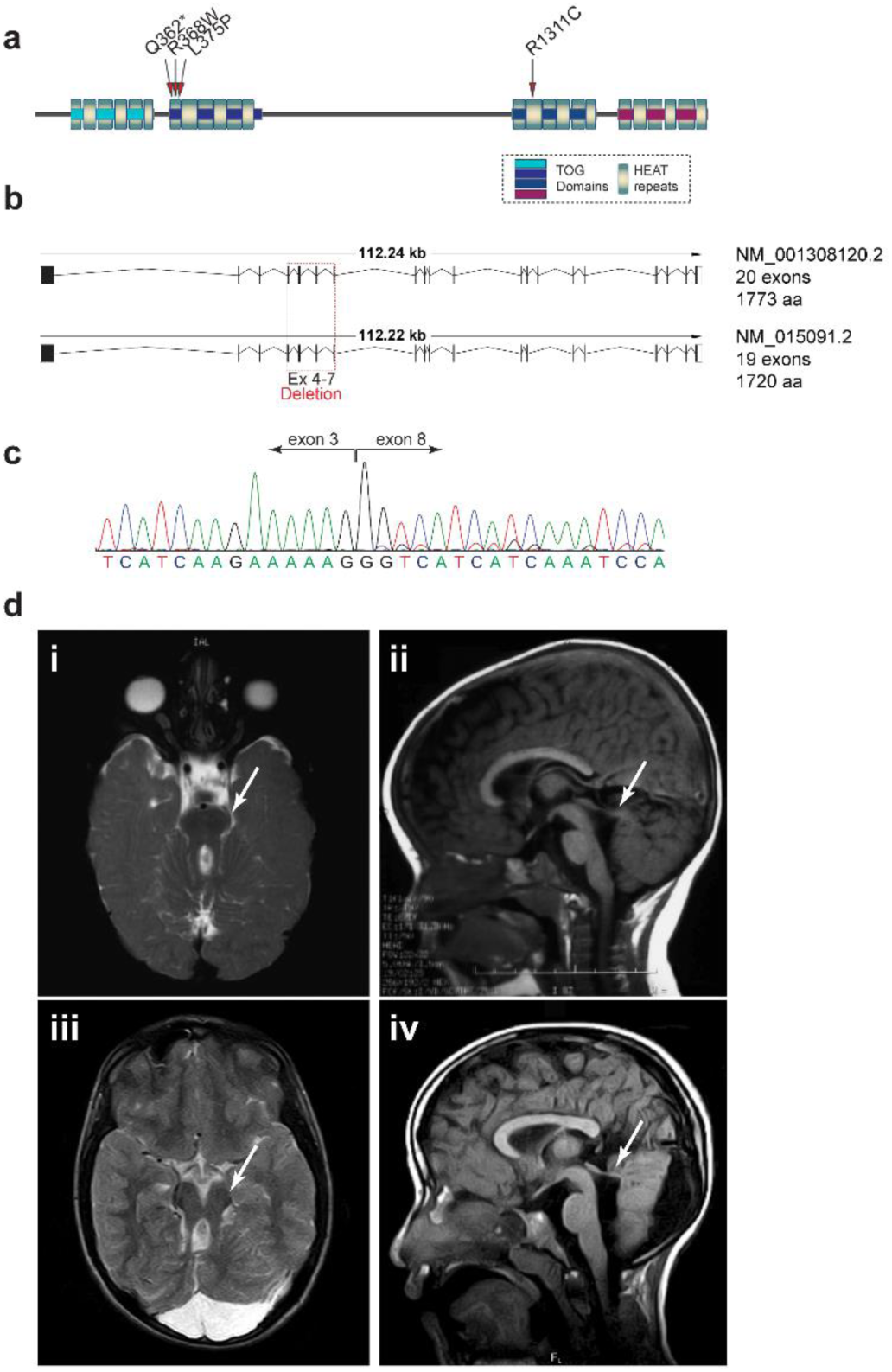
TOGARAM1 variants cause JBTS. **(A)** Protein schematic of TOGARAM1 with JBTS associated variants indicated, TOG domains 1-4 are shown with HEAT repeats (HR), HR are indicated in gradient blue. **(B)** Coding genomic schematic of *Homo sapiens* TOG array regulator of axonemal microtubules 1, TOGARAM1. Both transcript variant 1 (NM_001308120) and variant 2 (NM_015091.2) are shown. Patient UW360-3 exon 4-7 deletion is indicated. **(C)** Sanger sequencing results of the JBTS associated allelic deletion of exons 4-7. **(D)** Brain imaging features in individuals with *TOGARAM1*-related JBTS. Molar tooth sign, indicated by arrows in i and iii, and elevated roof of the 4th ventricle indicated by the arrows in ii and iv. The horizontally oriented superior cerebellar peduncles are visible in UW360-3 (i-ii) and 13DG1578 (iii-iv). The cerebellar tissue on the sagittal images is hemisphere based on the absence of vermis tissue on axial and coronal views (not shown). i and iii are sagittal T1-weighted images and ii and iv are axial T2- weighted images.

**Table 1.** Variants and Clinical Features in individuals with TOGARAM1-related JBTS

|  | **UW351-3** | **UW360-3** | **13DG1578** |
| --- | --- | --- | --- |
| **Variant 1\*** | c.1124T>C;<br>p.(Leu375Pro) | c.1084C>T; p.Gln362* | c.1102C>T; p.(Arg368Trp) |
| **Allele frequency 1 (33)** | 8/282484<br>2.83202E-05 | 0 | 0 |
| **CADD v1.3 (34)** | 25.5 | 35 | 28.7 |
| **Parent** | father | NA | NA |
| **Variant 2\*** | c.3931C>T;<br>p.(Arg1311Cys) | del14q21.2: g.45472226-45484228 | c.1102C>T; p.(Arg368Trp) |
| **Allele frequency 2** | 1/222730<br>4.48974E-06 | NA | 0 |
| **CADD v1.3<sup>3</sup>** | 35 | NA | 28.7 |
| **Parent** | mother | NA | NA |
| **Ethnicity/country** | Mixed European<br>(Australia) | Mixed European (US) | Middle Eastern (Egypt) |
| **Gender** | male | male | male |
| **Age** | 21 wk fetus | 16 yr | 11 yr, 5 mo |
| **Brain imaging** | vermis hypoplasia | MTS | MTS |
| **Dev disability** | NA | Y | Y |
| **Apnea/tachypnea** | NA | N | Y |
| **Abnormal eye movements** | NA | Y (and strabismus) | Y |
| **Retinal** | NA | N | N |
| **Kidney** | NA | N | N |
| **Liver** | NA | N | N |
| **Polydactyly** | Y | N | N |
| **Coloboma** | NA | N | N |
| **Craniofacial** | broad nasal bridge, post<br>rotated ears, thickened<br>neck | low set ears, high arched<br>palate | metopic ridge, frontal<br>bossing, deep set eyes,<br>hypertelorism, broad<br>nose, anteverted alae,<br>low set ears |
| **Other** | bilateral postaxial foot<br>polydactyly | widely spaced nipples,<br>undescended testes,<br>possible micropenis | widely spaced nipples,<br>small scrotum and testes |

\*NM\_015091.2
Abbreviations: mo=months, MTS=Molar Tooth Sign, N=No, NA=Not Available, US=United States, wk=weeks, Y=Yes, yr=years

The three affected individuals UW351-3, UW360-3, and 13DG1578 had features consistent with JBTS, including classic brain imaging findings (absent cerebellar vermis and thick, horizontally oriented superior cerebellar peduncles, giving the appearance of the molar tooth sign) in UW360-3 and 13DG1578 (Figure 2D; i-iv), and cerebellar vermis hypoplasia in fetus UW351-3. This fetus also had bilateral postaxial foot polydactyly and abnormal craniofacial features at autopsy including broad nasal bridge and posteriorly rotated ears. The two living children have typical hypotonia, ataxia, cognitive delays and behavioral features associated with JBTS, but no signs of retinal, kidney, or liver involvement. Uncommonly seen in individuals with JBTS, both boys were noted to have widely spaced nipples, genital abnormalities (undescended testicles and possible micropenis in UW360-3, and small scrotum and testicle in 13DG1578), and dysmorphic features (low-set ears in both with a high arched palate in UW360-3, and metopic ridge, frontal bossing, deep set eyes, and hypertelorism in 13DG1578).

### JBTS-associated *TOGARAM1* variants in the TOG2 domain disrupt the ARMC9-TOGARAM1 interaction

TOGARAM1 is a member of the highly conserved FAM179 protein family and is found across ciliated eukaryotes including *Chlamydomonas reinhardtii*, *Tetrahymena thermophila*, and *Caenorhabditis elegans*. TOGARAM1 has four conserved TOG domains that display similarity to the tubulin binding domains in ch-TOG and CLASP family proteins {Das:2015cv}. Arg368Trp and Leu375Pro lie within the highly conserved TOG2 domain (Figure 2A) which has been found to promote microtubule polymerization in vitro (26). The TOG2 domain conforms to the canonical TOG domain architecture found in other TOG array containing proteins such as CLASP and Stu2 (Supplemental Figure 2A, D), therefore disruption of this domain is predicted to disturb microtubule binding (35). Using in silico analysis of the effects of point mutations, we found that both Arg368Trp and Leu375Pro mutations were predicted to be deleterious to protein structure (35). While residue 368 is not highly conserved, modeling the wild-type and mutant TOG domains using HOPE (35) predicts that the larger and neutral tryptophan disrupts the normal hydrogen bonds with the aspartic acid residues at positions 361 and 405 (Supplemental Figure 2B, C). Position 375 is located in a predicted α-helix, and is also not highly conserved; however, proline introduces a bend in the polypeptide chain different from other amino acids and the substitution of a proline is predicted to disrupt this α-helix (Supplemental Figure 2E, F), likely affecting protein folding or interaction with other domains (35).

To assess the effects of JBTS-associated variants, we expressed wild-type and mutant mRFP-tagged TOGARAM1 (Arg368Trp, Leu375Pro, and Arg1311Cys) in control (Figure 3B-E) and in a genetically edited *TOGARAM1* mutant hTERT-RPE line (Supplemental Figure 3A-B). The genetically edited *TOGARAM1* mutant hTERT-RPE line, *TOGARAM1* mut 1, has a biallelic deletion of the ATG site of TOGARAM1 (Supplemental Figure 3C; Supplemental Figure 4A, B). Overexpressed wild-type TOGARAM1 localized along the ciliary axoneme and was associated with markedly longer cilia compared to untransfected wild-type cells (Figure 3A, B) and *TOGARAM1* mut 1 cells (Supplemental Figure 3A,B). All three variants also localized to the cilium, but overexpression of TOG2-domain variants Arg368Trp and Leu375Pro resulted in longer cilia while overexpression of the TOG3-domain variant Arg1311Cys resulted in shorter cilia compared to untransfected cells (Figure 3A-E, quantification in Figure 3F, Supplemental Figure 3A,B). These data suggest that disruption of the TOG3 domain may have a dominant negative effect on the microtubule polymerization capacity of TOGARAM1 and disruption of the TOG3 domain has a different effect on TOGARAM1 localization and ciliary extension as compared to TOG2 domain variants.

**Figure 3.**
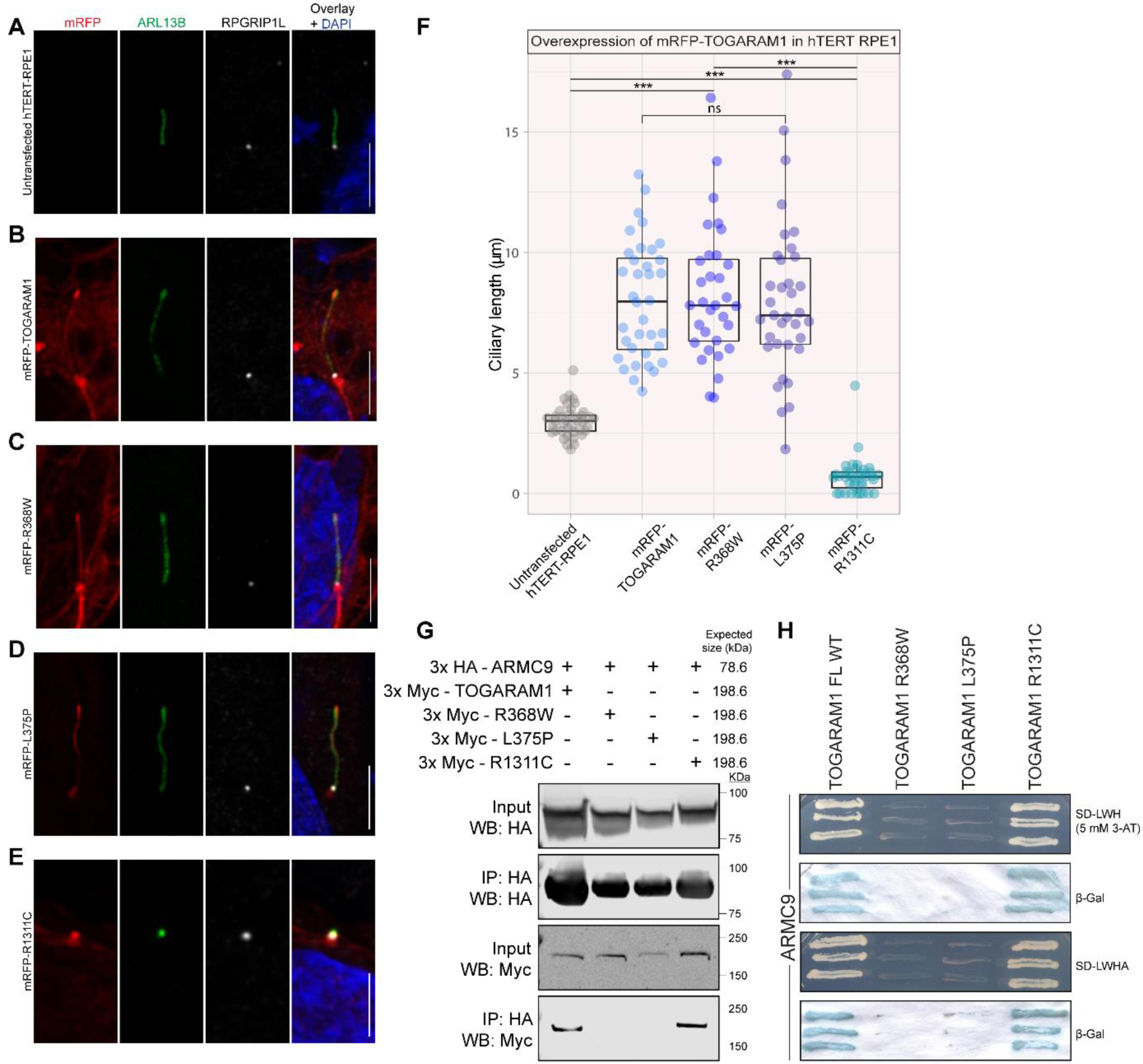
JBTS variants cause altered ciliary length and disruptions in TOG2 abrogate the interaction with ARMC9. **(A)** Untransfected control hTERT-RPE1 cells showing ciliary length in the absence of transfection, the cilium is shown with transition zone marker RPGRIP1L (white) and ciliary membrane marker ARL13B (green). **(B)** Transient overexpression in hTERT-RPE1 cells of mRFP-TOGARAM1 WT (red), **(C)** mRFP- Arg368Trp TOGARAM1 (red), **(D)** mRFP-Leu375Pro TOGARAM1 (red), and **(E)** mRFP-Arg1311Cys TOGARAM1 (red) shown with transition zone marker RPGRIP1L (white) and ciliary membrane marker ARL13B (green). Images are representative of >30 cilia assessed per condition. Scale bars for a-e are 5 μm. **(F)** The length of >30 cilia measured from cells overexpressing mRFP-TOGARAM1 or variants. 39 cilia from untransfected cells were measured for comparison (n=36 cilia for mRFP-TOGARAM1 WT, n=32 cilia for mRFP-Arg368Trp, n=35 cilia for mRFP-Leu375Pro and n=36 cilia for mRFP-Arg1311Cys). p<0.0001 between cilia from untransfected cells and mRFP-TOGARAM1, mRFP-Arg368Trp, mRFP-Leu375Pro, and mRFP- Arg1311Cys. No significant differences were found between the ciliary lengths of cells overexpressing mRFP-TOGARAM1, mRFP-Arg368Trp, and mRFP-Leu375Pro. The difference between mRFP-Arg1311Cys and mRFP-TOGARAM1, mRFP-Arg368Trp, and mRFP-Leu375Pro was significant (p<0.0001) by one-way ANOVA and Tukey’s multiple comparison test. Box plot horizontal bars represent the median. **(G)** Co-immunoprecipitation of HA-tagged ARMC9 and Myc-tagged TOGARAM1: Wild-type and Myc-tagged Arg1311Cys interact with ARMC9 while TOGARAM1 variants Arg368Trp and Leu375Pro do not, indicating a loss of interaction due to variants in the TOG2 domain. **(H)** A Y2H direct interaction analysis assay with ARMC9 and TOGARAM1: Wild-type and Arg1311Cys TOGARAM1 interact with ARMC9 while the TOGARAM1 variants Arg368Trp and Leu375Pro do not, again indicating a loss of interaction due to TOG2 variants.

We next investigated the effects of the *TOGARAM1* variants on the interactions with ARMC9 using co-IP, binary yeast two-hybrid (Y2H) interaction analysis, and PalmMyr colocalization assays. We found that the variants in the TOG2 domain (Arg368Trp and Leu375Pro) abolished co-IP of HA-ARMC9 with Myc-TOGARAM1, while the Arg1311Cys variant, occurring in the TOG3 domain, appears not to influence this interaction (Figure 3G). Y2H analysis confirmed these binary interactions (Figure 3H). In PalmMyr assays, the individually-expressed wild-type and mutant mRFP-tagged TOGARAM1 proteins localized along the length of cilia and along cytoplasmic microtubules (Supplemental Figure 3C). Co-expression of PalmMyr-CFP tagged ARMC9 with wild-type or Arg1311Cys mRFP-TOGARAM1 resulted in strong co-localization along microtubules (Supplemental Figure 3D), indicating protein-protein interaction. In contrast, co-expression with the TOG2 variants resulted in PalmMyr-CFP tagged ARMC9 remaining localized to the plasma membrane, suggesting a lack of interaction with the TOGARAM1 mutants affecting the TOG2 domain (Supplemental Figure 3D). Taken together, these data indicate that variants in the TOG2 domain abrogate the ARMC9 - TOGARAM1 interaction.

### *togaram1* mutations cause ciliopathy phenotypes in zebrafish

To further investigate the function of TOGARAM1 and the association between TOGARAM1 dysfunction and JBTS, we turned to zebrafish, an established model organism for ciliopathies. Indeed, zebrafish display a wide variety of ciliated cell types similar to humans and mutations in human ciliopathy genes result in typical ciliopathy phenotypes. Ciliated cells typically assessed in the zebrafish model include epithelial cells in pronephric (kidney) ducts, olfactory neurons in nose pits, neuronal progenitors on larval brain ventricular surfaces or cells lining the Kuppfer’s vesicle (equivalent to the mouse embryonic node). We identified a single zebrafish *togaram1* ortholog, which is not described in Ensembl and incompletely annotated in NCBI. Zebrafish *togaram1* displays a highly conserved C-terminal region encompassing two TOG domains (similar to human TOG3 and 4) and a single N-terminal TOG domain corresponding to mammalian TOG2 domain (Supplemental Figure 5A). While the linker region between TOG2 and TOG3 is poorly conserved, the three TOG domains are well-conserved between zebrafish and its corresponding human counterparts (50-58% amino acid identity and 72-77% similarity). Gene synteny analysis confirmed that the identified zebrafish sequence represents the ortholog of human *TOGARAM1* (Supplemental Figure 5B). Importantly, on the paralogous chromosomal fragment generated by the teleost-specific whole genome duplication, no second *togaram1* paralog could be identified. Moreover, synteny analysis also revealed that the zebrafish genome lacks a *TOGARAM2* ortholog (Supplemental Figure 5C), leaving zebrafish with just one *togaram* ortholog. These findings support the utility of zebrafish as a model for *TOGARAM1*-associated human disease.

We next generated zebrafish mutants using CRISPR/Cas9. Two different pairs of sgRNAs targeting different regions of the gene (Supplemental Figure 5D) led to similar phenotypes in injected F0 larvae. 39% developed a curved body shape and 9% developed kidney cysts, both typical zebrafish ciliopathy-associated phenotypes (Supplemental Figure 5E). Single larvae were lysed for genotyping which revealed a very high mutation efficiency (94% of sequenced clones from 7 larvae had small insertions-deletions, the majority of which were frameshift mutations). Among the mutant F0 fish, a striking scoliosis phenotype was observed in juveniles (Supplemental Figure 5E), reminiscent of other ciliopathy mutants including *armc9* CRISPR-F0 fish (24). Taken together, these results confirm that loss of *togaram1* causes ciliopathy phenotypes in zebrafish and supports a role for *togaram1* in ciliary function.

### *armc9* and *togaram1* mutant zebrafish display similar phenotypes

To further evaluate the link between *TOGARAM1* and *ARMC9*, we compared zebrafish mutants in the two genes. Following up on our previous work (24), we raised several stable (>F2) zebrafish lines harboring frameshift insertion and deletion alleles of *armc9* (Supplemental Figure 5F). Since homozygous mutants from all generated alleles have comparable phenotypes, we focused on the *armc9*^zh505^ allele harboring a 110bp insertion in exon 14 for follow up experiments (Supplemental Figure 5F). *armc9*-/- larvae have a straight body shape and an incompletely penetrant pronephric cyst phenotype affecting 44% of homozygous mutants (Figure 4B). In comparison, *togaram1 ^zh509^ ^or^ ^zh510^* mutant F2 larvae harboring frameshift mutations in exon 20/21 have a slightly curved body shape and display a similar rate of kidney cysts compared to *armc9*-/- mutants (Figure 4C). Pronephric cysts and body curvature do not necessarily correlate with each other in *togaram1* mutants, as each phenotype can be found in isolation or in combination, but overall, 85 % of *togaram1*-/- larvae have at least one ciliopathy phenotype. In addition to frameshift mutations in exon 20/21, we also identified a 21 bp in-frame deletion in the TOG4 domain leading to loss of 6 highly conserved amino acids (*togaram1^zh508^*, Supplemental Figure 6). Homozygous in-frame mutant larvae are indistinguishable from the frameshift mutants, suggesting that the TOG4 domain may be critical for Togaram1 function.

**Figure 4.**
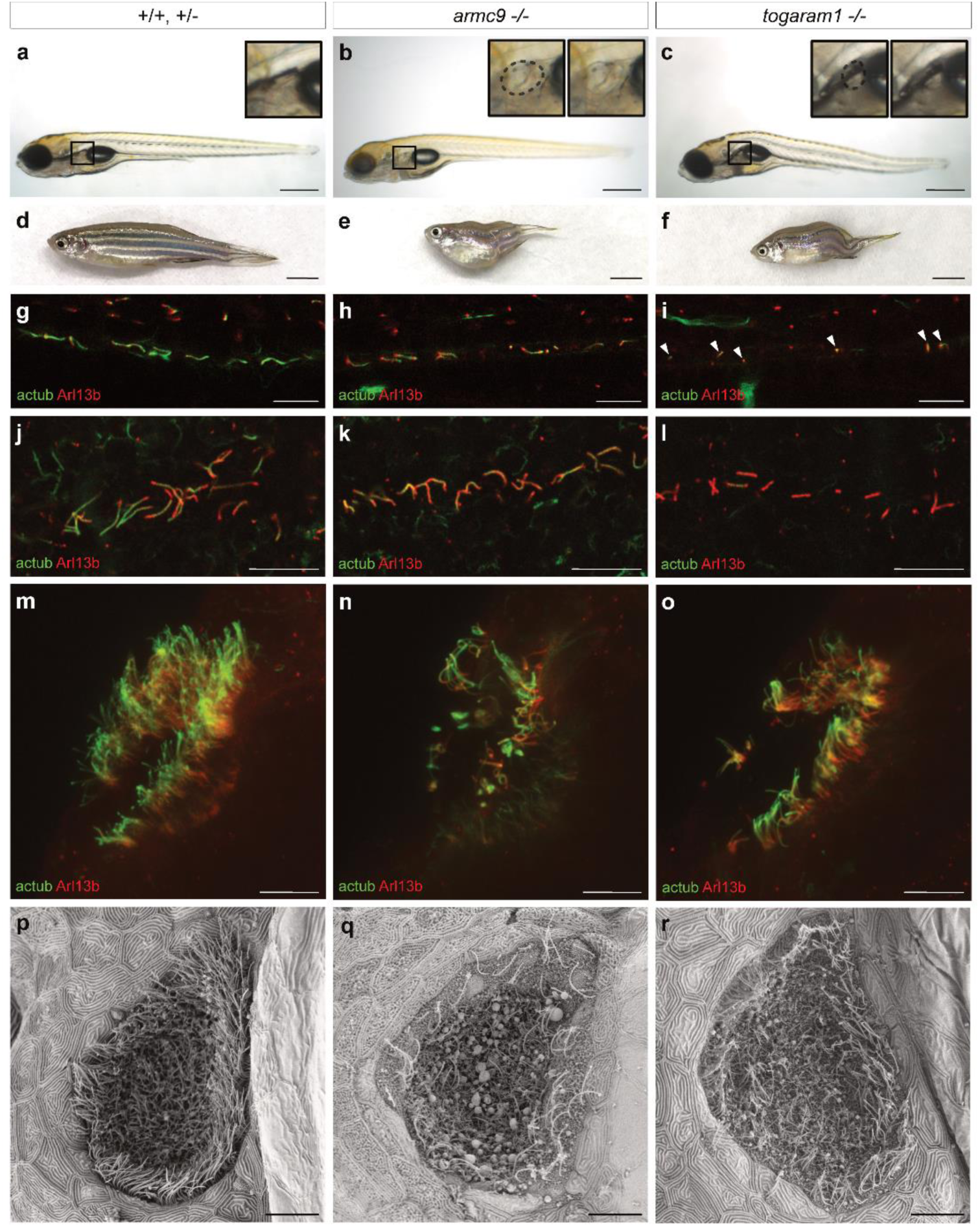
*armc9* and *togaram1* mutant zebrafish display ciliopathy associated phenotypes. **(A-C)** Larval phenotype demonstrating kidney cysts in *armc9*-/- (B) and kidney cysts and slightly curved body shape in *togaram1*-/- (C). Black boxes in (A-C) show magnification of glomerulus region in inset. Dashed lines highlight kidney cysts in (B, C). **(D-F)** Adult scoliosis phenotype in both mutants (E, F) compared to wild-type (D). **(G-I)** Immunofluorescence of midbrain ventricles shows shortened cilia in 3 day post fertilization (dpf) *armc9* and *togaram1* mutant zebrafish larvae (H, I). **(J-L)** Immunofluorescence of 3 dpf zebrafish nosepits: decreased cilia number in both mutants (K, L) compared to wildtype (J). **(M-0)** Scanning electron microscopy of 5 dpf zebrafish nosepits confirming reduced cilia numbers in *armc9*-/- (N) and *togaram1*-/- (O). Scale bars are 500 μm in (A-C), 5 mm in (D-F) and 10 μm in (G-O).

In addition to the proneprhic cyst phenotypes observed, both *armc9-/-* and *togaram1*-/- fish develop scoliosis as juveniles (Figure 4A, F), as previously described in other zebrafish ciliary mutants Given that both pronephric cysts and curved bodies are typical ciliopathy phenotypes, we next analyzed the cilia in both mutants using immunofluorescence staining with anti-Arl13b and anti-acetylated α-tubulin antibodies. We observed reduced numbers of shortened larval pronephric, ventricular (Figure 4H, I) and nose pit (Figure 4K, L) cilia in both mutants compared to wild-type (Figure 4G, J), the latter being confirmed by scanning electron microscopy (SEM) (Figure 4M-O). The reduced and shortened cilia in both mutants support a role for Armc9 and Togaram1 in zebrafish cilium formation and/or stability.

### JBTS-associated ARMC9 and TOGARAM1 variants result in decreased ciliary length

To gain insight into the ciliary defects associated with JBTS in humans, we evaluated four fibroblast lines from patients with *ARMC9*-associated JBTS. Western blot analysis of total protein lysates revealed that all four cell lines express trace levels of the two major ARMC9 isoforms at 92kDa and 75.5kDa seen in control fibroblasts (Figure 5A, B). To evaluate for ciliary effects, we serum starved control and ARMC9 mutant cells and stained with anti-acetylated α-tubulin and anti-ARL13B. All four patient lines displayed significantly shorter cilia, with an average length of 2.3± a standard deviation (SD) of 0.7 −3.3± SD of 1.4 μm compared to an average length of 3.6± SD of 1.4 μm in controls (Figure 5C). When assessing ciliation levels, we found that approximately 80% of control cells had cilia after 48 hours without serum, compared to 75-86% in the patient cell lines (Figure 5D), suggesting that ARMC9 does not play an integral role in ciliogenesis.

**Figure 5.**
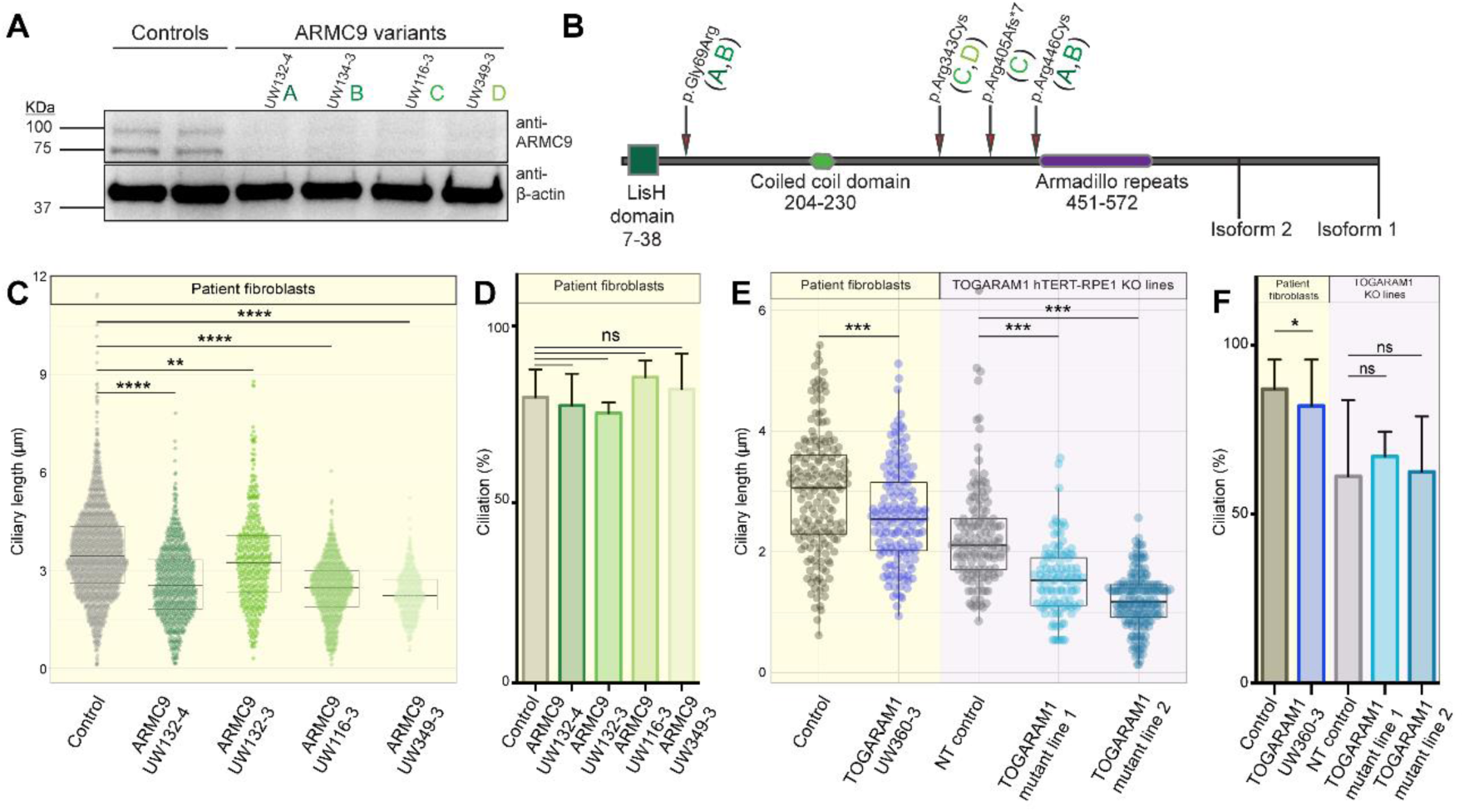
*ARMC9* and *TOGARAM1* variants result in short cilia. **(A)** Immunoblot of endogenous ARMC9 in control and patient fibroblasts indicating trace amounts of ARMC9 isoform 1 and 2 (92 and 75.5 kDa) in patient fibroblasts. Loading control is β-actin. **(B)** ARMC9 schematic indicating JBTS-associated patient variants (green letters represent variants found in patients as indicated in a). Isoform 1 is 818 amino acids, and isoform 2 is 665 amino acids. **(C)** Ciliary length measurements in control and *ARMC9* patient fibroblasts (control = 1395 cilia, UW132-4 = 699 cilia, UW132-3 = 437 cilia, UW116-3=656 cilia, and UW349-3=353 cilia). p<0.0001 between control and *ARMC9* patient cilia by one-way ANOVA. **(D)** *ARMC9* fibroblast ciliation levels (control= 1723 cilia, UW132-4 = 898 cilia, UW132-3 = 584 cilia, UW116- 3= 764 cilia, and UW349-3 = 425 cilia). Results were not significantly different per one-way ANOVA. **(E)** Ciliary length measurements in control and *TOGARAM1* patient fibroblast line (yellow panel). p=0.0003 per unpaired t-test. hTERT-RPE1 cilia length in wild-type and *TOGARAM1* mut lines (purple panel) based on ARL13B ciliary staining. >100 cilia were pooled from 2 independent experiments (n=137 cilia for wild type, n=111 cilia for *TOGARAM1* mut line 1, and n=178 cilia for *TOGARAM1* mut line 2. p<0.0001 per one-way ANOVA. Box plot horizontal bars represent the median. **(F)** *TOGARAM1* fibroblast ciliation levels >400 cells were pooled from 3 experiments (n= 466 for control line and n= 429 for UW360-3). p=0.0164 per unpaired t-test. Measurements of hTERT-RPE1 ciliation percentages in wild-type and *TOGARAM1* mut: >330 cells were pooled from 2 independent experiments (n= 330 cells for wild type, n= 363 cells for TOGARAM1 mut line 1, and n = 357 cells for TOGARAM1 mut line 2). Results were not significantly different per one-way ANOVA. Symbols indicate the following P values: ns, P > 0.05; *, P ≤ 0.05; **, P ≤ 0.01; ***, P ≤ 0.001; ****, P ≤ 0.0001.

The single fibroblast line from patient UW360-3 with *TOGARAM1*-associated JBTS had a slightly lower ciliation rate than control (85% versus 91% respectively) (Figure 5F). Mean ciliary length was also shorter (2.61 ± SD of 0.81 μm, n = 154 cilia) versus control (2.97 ± SD of 0.99 μm, n = 179 cilia) (Figure 5E). To generate additional data about the effects of loss of *TOGARAM1* function on cilia in mammalian cells, we turned to CRISPR/Cas9 genome-edited *TOGARAM1* hTERT-RPE1 mutant cells. gRNAs targeting the translation start site of exon 1 resulted in two different lines harboring a disruption in the ATG site of both alleles of *TOGARAM1* (Supplemental Figure 4A, B). These lines make significantly shorter cilia (mut line 1: average ciliary length = 1.54 ± a SD of 0.59 μm, n= 111 cilia; and mut line 2: average ciliary length = 1.18 ± a SD of 0.46 μm, n = 178 cilia) compared to the isogenic control (average ciliary length = 2.27 ± a SD of 0.84 μm, n = 137 cilia) (Figure 5E). Ciliation levels in the TOGARAM1 mutant/engineered lines did not differ significantly from the isogenic control (Figure 5F). Taken together these results suggest that disruptions in *TOGARAM1* and *ARMC9* lead to shorter ciliary length but do not affect overall ciliation rates.

### Transition zone integrity with ARMC9 and TOGARAM1 dysfunction

Given the well-described role of the transition zone (TZ) in JBTS (38), we sought to examine the integrity of this compartment with TOGARAM1 or ARMC9 dysfunction. Since TZ dysfunction often results in loss of ciliary ARL13B, which secondarily causes loss of ciliary INPP5E (39, 40), we performed quantitative immunofluorescence (qIF) on control, *ARMC9*, and *TOGARAM1* patient cell lines. Our data revealed mildly decreased levels of ARL13B in three of four *ARMC9* patient fibroblast lines and normal levels in the *TOGARAM1* patient fibroblast line (Figure 6A; Supplemental Figure 7A, C). Importantly, the mild ARL13B decrease observed in 3 of 4 ARMC9 lines was not associated with a concomitant loss of ciliary INPP5E (Figure 6B; Supplemental Figure 7B), indicating that lower ARL13B levels were still sufficient to properly localize INPP5E. In zebrafish, Arl13b levels were not lower in either mutant (and even slightly increased in the *armc9*-/- fish) (Figure 6A, E, F). Together, these results are strikingly different and much less severe than the marked ciliary ARL13B and INPP5E reduction observed in TZ mutants (39, 40). To evaluate the composition of the TZ directly, we performed immunostaining for canonical TZ proteins RPGRIP1L in human cell lines (Figure 6G, H) and Cc2d2a in zebrafish (Figure 6F). Both proteins localized normally to the TZ of the respective cilia. Taken together, these findings suggest that the TZ is generally intact, despite dysfunction of the ARMC9-TOGARAM1 complex.

**Figure 6.**
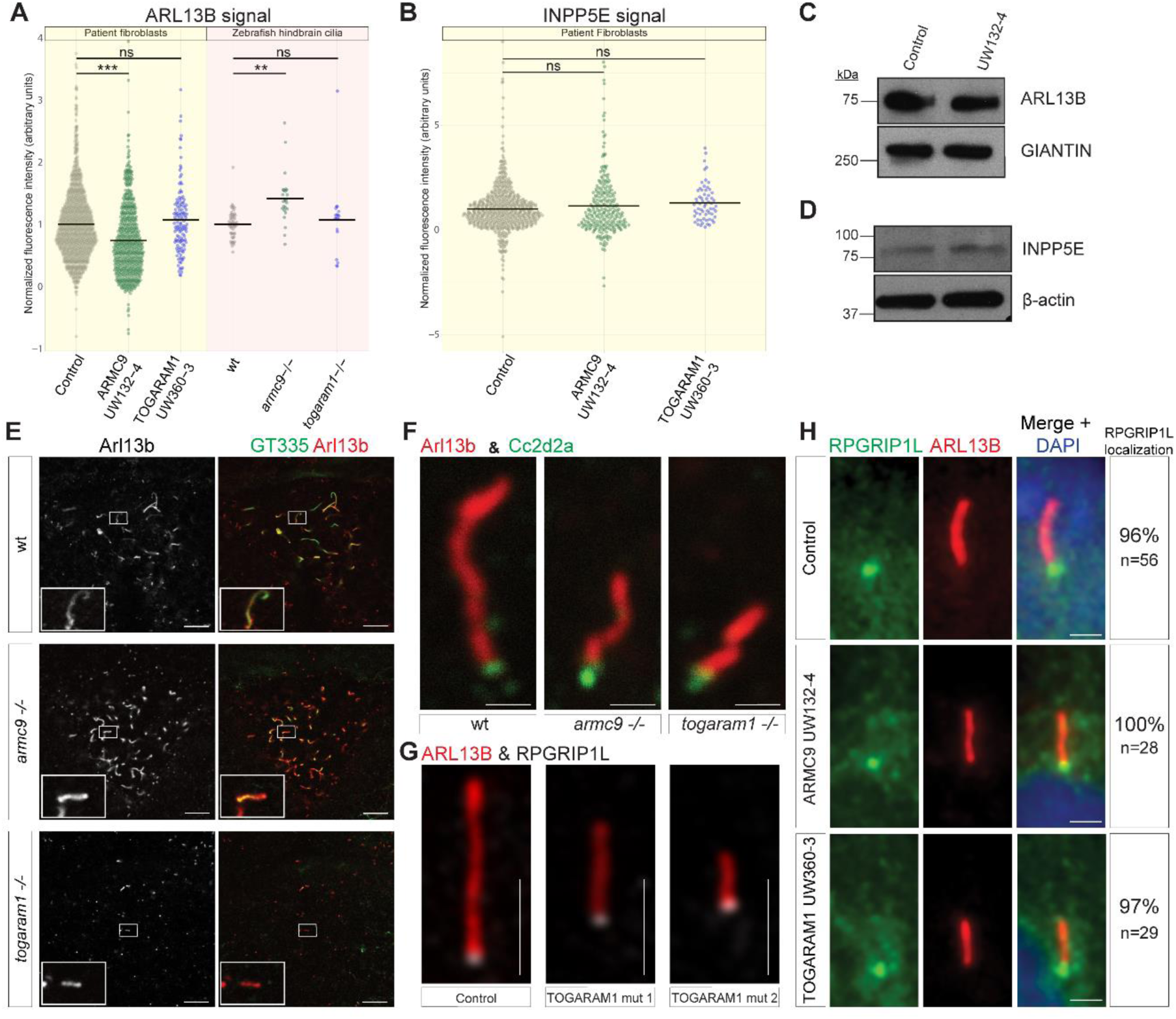
ARMC9 or TOGARAM1 dysfunction does not grossly affect the transition zone. **(A)** Relative fluorescence intensity of ARL13B signal in control (grey), *ARMC9* (green) and *TOGARAM1* (blue) in patient fibroblasts (yellow panel) and in zebrafish hindbrain cilia (pink panel). All controls are normalized to 1. Bars represent the mean. For patient fibroblasts >120 cells were pooled and analyzed (n= 1089 control line, n=582 UW132-4 and n= 126 UW360-3). For KO zebrafish, 10 cilia per larva were measured (n=56 wt larvae, n=24 *armc9* -/-larvae, and n=19 *togaram1* -/- larvae; data pooled from three independent experiments). **(B)** Relative fluorescence intensity of INPP5E signal in control (grey), *ARMC9* (green) and *TOGARAM1* (blue) patient fibroblasts. >60 cells were pooled and analyzed (n= 620 control line, n=248 UW132-4 and n= 62 UW360-3). Signal intensity of ARL13B and INPP5E across all ARMC9 patient fibroblast lines can be found in Supplemental Figure 7. P values: **(C)** Western blotting analysis of ARL13B **(D)** and INPP5E in *ARMC9* UW132-4 patient fibroblasts. GIANTIN and β-actin serve as loading controls respectively. **(E)** Representative immunofluorescence Arl13b (red) and GT335 (green) staining of 3 dpf zebrafish hindbrain cilia (single confocal section). Left panels show only Arl13b signal, insets are magnified single cilia. Scale bars 10 μm. Quantification of fluorescence intensity is shown in the pink panel in (A). **(F)** Single hindbrain cilium stained with Arl13b (red) and Cc2d2a (green) in 3dpf control, *armc9-/-* and *togaram1-/-* zebrafish. Scale bars 1 μm. **(G)** Immunofluorescence image of RPGRIP1L (white) and ARL13B (red) localization in cilia from WT hTERT-RPE1 and two *TOGARAM1* mutant lines. Scale bars 2 μm. **(H)** RPGRIP1L localization in *ARMC9* and *TOGARAM1* patient fibroblasts. Percentage of cilia with robust RPGRIP1L puncta as indicated in figure. Scale bars 2 μm. Images are representative of all cilia assessed. All p-values were determined by ordinary one-way ANOVA; n s P > 0.05; ** P ≤ 0.01; *** P ≤ 0.001.

### ARMC9 and TOGARAM1 dysfunction affects tubulin post-translational modifications in patient fibroblasts and zebrafish mutants

When measuring ciliary ARL13B and INPP5E, we noted that the acetylated α-tubulin and polyglutamylated signal appeared substantially less intense in patient cell lines versus controls (Fig 7A, B). Using qIF, we found that mean acetylated tubulin levels were ∼50% of control levels in the *ARMC9* lines and ∼70% of controls in the *TOGARAM1* line (Figure 7E, F and Supplemental Figure 8A, B). We found that the mean polyglutamylated signal levels were ∼35% and ∼45% of control levels in ARMC9 and TOGARAM1 lines respectively (Fig 7B, F and Supplemental Fig 7). Western blots of whole cell lysates also demonstrated substantially lower levels of both acetylated and polyglutamylated tubulin in *ARMC9* fibroblast lines compared to controls (Figure 7A, B). We observed similar reductions of ciliary acetylated and polyglutamylated tubulin in the remaining ventricular cilia of zebrafish with *armc9* and *togaram1* mutations (Figure 7C-F). Together, these results indicate that loss of either ARMC9 or TOGARAM1 results in decreased post-translational modifications of axonemal tubulin across multiple model systems.

**Figure 7.**
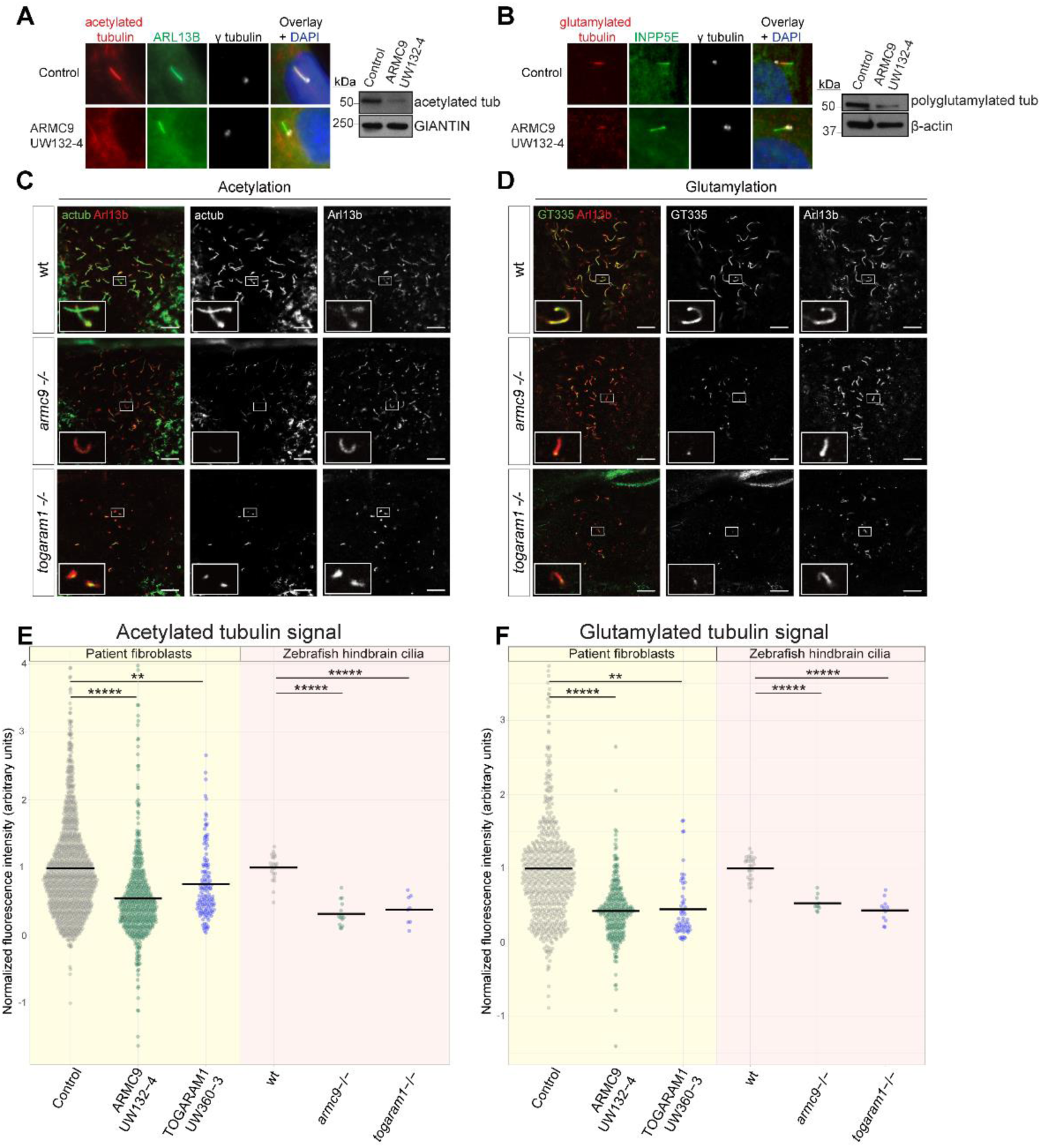
ARMC9 and TOGARAM1 mutant cilia display reduced tubulin posttranslational modifications in both patient fibroblasts and zebrafish ventricular cells. **(A–B)** IF and immunoblots of (A) acetylated and (B) polyglutamylated tubulin in *ARMC9* patient fibroblasts versus control. In the immunoblots, GIANTIN and β-actin are loading controls. **(C)** Whole mount immunofluorescence of 3 dpf hindbrain cilia in zebrafish, stained with anti-Arl13b (red) and anti-acetylated α-tubulin (green). **(D)** 3 dpf hindbrain cilia in zebrafish stained with anti-Arl13b (red) and anti-GT335 (green). Scale bars represent 10 μm. Note that anti-acetylated tubulin also stains axons in the developing brain, visible at the edges of the image in **(C)**. **(E)** Quantification of acetylated tubulin signal from patient fibroblasts (yellow panel) n= 1106 cilia for control line, n=532 cilia for UW132-4 and n= 131 cilia for UW360-3; in pink panel, for KO zebrafish, 10 cilia per larva were analyzed, n=23 wt larvae, n=14 *armc9* -/- larvae, and n=8 *togaram1* -/- larvae; data pooled from three independent experiments. **(F)** Polyglutamylation signal assessed in n= 602 cilia for control line, n=298 cilia for UW132-4 and n= 58 cilia for UW360-3; for KO zebrafish n=28 wt larvae, n=10 *armc9* -/- larvae, and n=11 *togaram1* -/- larvae. Quantification of signal was normalized to wt, zebrafish mutants are shown in the pink panel and patient-derived fibroblasts are shown in the yellow panel of the quantification graphs. *ARMC9* patient line UW132-4 is representative of all *ARMC9* patient lines assessed. In (E and F), data points above 4 and below −2 have been excluded from this graph for display clarity, but these data points were retained in the statistical analysis. For a complete graph of all data points and for a graphical summary of all *ARMC9* lines refer to Supplemental Figure 8 (A, B) and (C, D) respectively. All p-values were determined as significant by one-way ANOVA.

### ARMC9 and TOGARAM1 dysfunction is associated with abnormal ciliary resorption

Post-translational modifications of microtubules such as acetylation and polyglutamylation are enriched in the ciliary compartment and play roles in ciliogenesis, axoneme stability, and cilium disassembly (13). To investigate the consequence of reduced ciliary microtubule post-translational modifications on axonemal stability, we evaluated cilia of control and *ARMC9* mutant cells for sensitivity to cold-induced microtubule depolymerization. In control cells, a 10-minute treatment at 4°C was not associated with reduced numbers of cilia, while cold-treated *ARMC9* patient cells had 20-40% fewer cilia than untreated cells (Figure 8A). *TOGARAM1* patient cell cilia were also more susceptible to cold-induced depolymerization, with 20% fewer cilia after treatment compared to untreated cells (Fig 8A).

**Figure 8.**
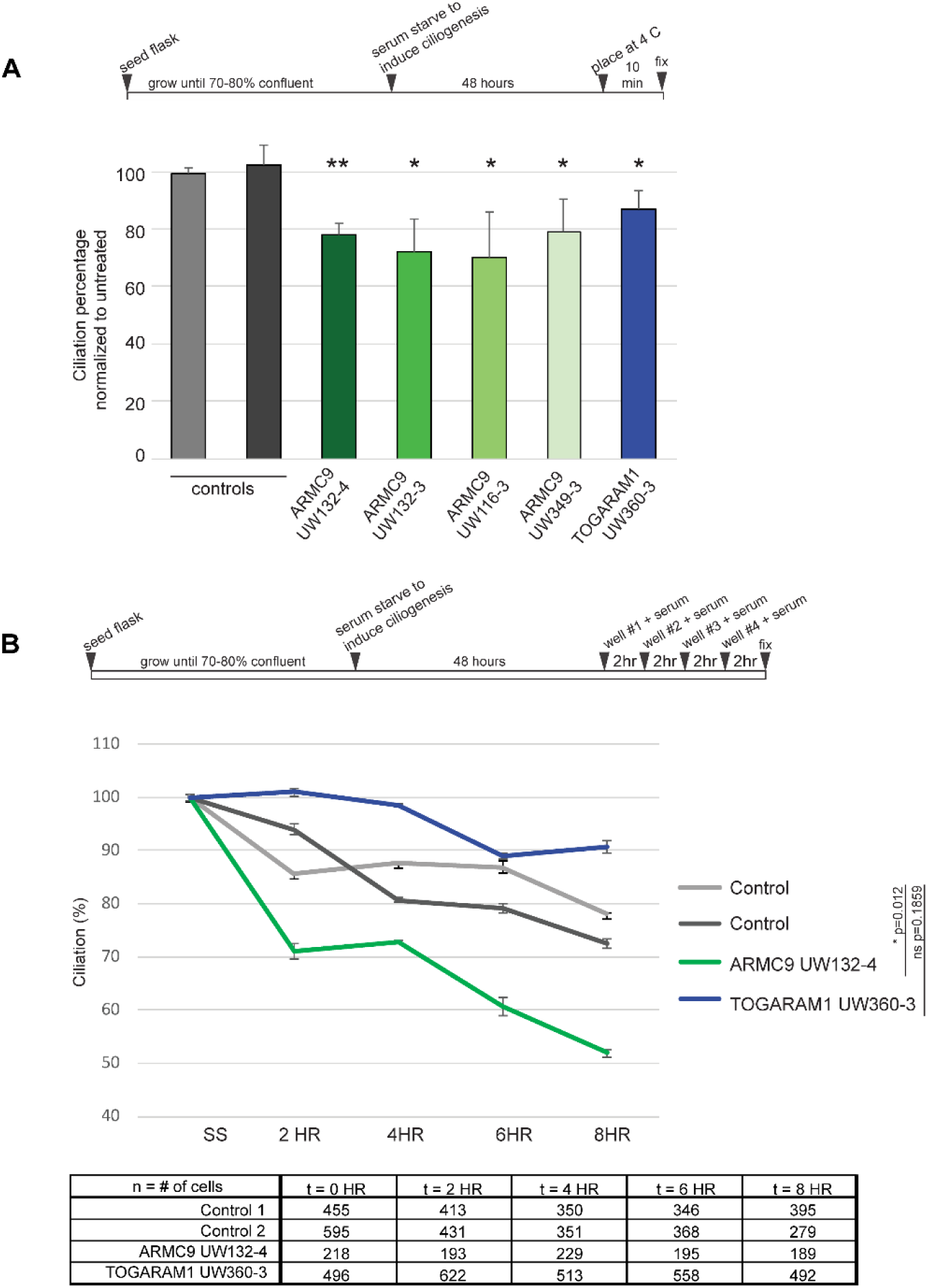
JTBS patient fibroblasts exhibit abnormal axonemal stability. **(A)** Cold-induced depolymerization assay schematic and ciliation percentages of treated cells normalized to non-treated controls. **(B)** Assessment of ciliation levels 2, 4, 6, and 8 hours post serum add back to fibroblasts starved for 48 hours, >180 cells were assessed for each condition. For exact n values refer to the table beneath the bar graph. Ciliation percentages were normalized to 100% at the time of serum resupplementation, percentages represent the amount of remaining cilia compared to the time of resupplementation.

As a second measure of cilium stability, we evaluated the rate of cilium resorption after serum re-addition to serum-starved cells. Serum provides growth factors that quickly initiate ciliary resorption, so that cells can re-enter the cell cycle. In controls, the ciliation rate was ∼85% of baseline 4 hours after serum re-addition. In contrast, the ciliation rate was 70% of baseline in *ARMC9* patient fibroblasts only 2 hours after serum re-addition, and by 8 hours, it was down to 50%, compared to 75% in controls (Figure 8B). To determine whether the faster resorption was due to an overactive deacetylating enzyme in the ARMC9 cell lines, we repeated these experiments with and without the histone deacetylase 6 (HDAC6) inhibitor tubacin. Tubacin treatment did not rescue the faster resorption in ARMC9 cell lines to control levels (Supplemental Figure 9A-C). Intriguingly, the ciliation rate of the one *TOGARAM1* patient fibroblast line available remained 90% of baseline even 8 hours after serum re-addition, substantially higher than controls (Figure 8B). These data suggest that the ARMC9-TOGARAM1 complex plays a role in regulation of axonemal stability.

## Discussion

In this study, we identified a new JBTS-associated protein module that can be distinguished physically and functionally from the previously proposed JBTS protein network at the transition zone of primary cilia (41). Several components of this new module localize at the ciliary basal body (24) and at the proximal end of the ciliary axoneme (25, 42). Mutations in the genes encoding two directly interacting members of the module, ARMC9 and TOGARAM1, result in defects in cilium length, microtubule post-translational modifications (acetylation and polyglutamylation), and ciliary stability in patient-derived fibroblasts, zebrafish mutants, and genetically edited hTERT-RPE1 cell lines (Summary Figure 9).

**Figure 9.**
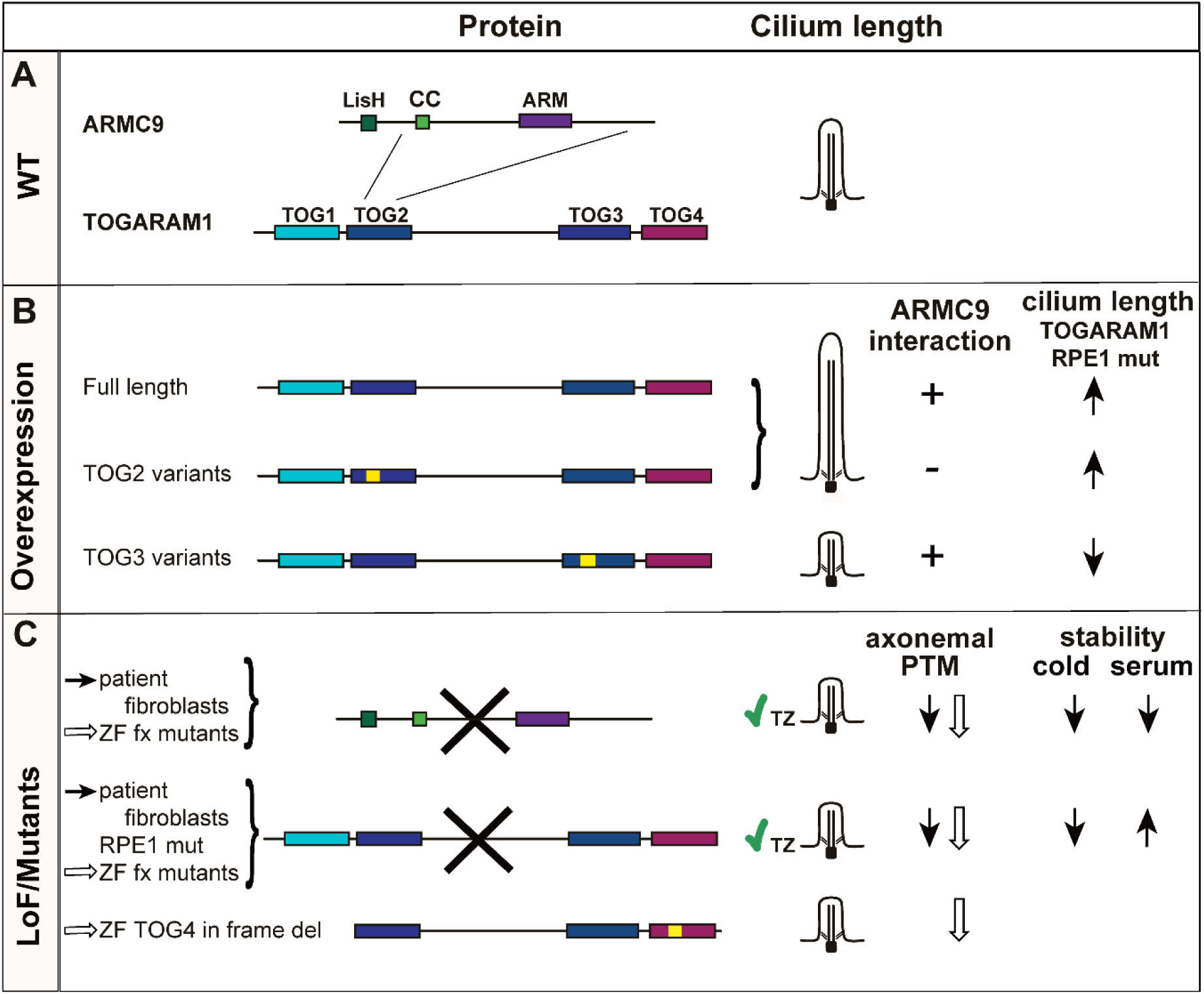
Graphical Summary: Disruptions of the ARMC9-TOGARAM1 complex affect ciliary length and resorption. **(A)** TOGARAM1 interacts with ARMC9 through its TOG2 domain. **(B)** Effects of overexpression of *TOGARAM1* wild-type and JBTS-associated variants on ciliary length in RPE1 *TOGARAM1* mut cells and consequences of these variants on the interaction with ARMC9. **(C)** Consequences of mutations in *ARMC9* or *TOGARAM1* on ciliary length and axonemal post-translational microtubule modifications (PTM) in patient fibroblast lines (black arrows) or zebrafish mutants (white arrows). Transition zone (TZ) integrity despite mutations in complex members is indicated with a green checkmark. Consequences of *TOGARAM1* and *ARMC9* mutations on ciliary resorption induced by cold or by serum re-addition in patient fibroblasts are indicated with black arrows. Yellow boxes represent mutations. Bold crosses indicate presumed loss-of-function mutations. *del deletion, fx frameshift, LoF loss of function, WT wild-type, ZF zebrafish. RPE1 mut hTERT-RPE1 TOGARAM1 mutant lines. Protein domains: LisH Lis-homology, CC coiled-coil, ARM armadillo, TOG tumor overexpression gene*.

### The ARMC9-TOGARAM1 complex in JBTS

Knowledge of the components and associations of the ciliary molecular machines has been instrumental to acquiring insights into the correlation of ciliopathy genotypes, a wide spectrum of overlapping phenotypes, and the associated pathomechanisms. Several affinity and proximity proteomics approaches have been used to determine the topology of ciliary protein-protein interaction networks and generate molecular blueprints of the ciliary machinery, e.g. the entire ciliary organelle (43), of the human centrosome-cilium interface (42), or specific ciliopathy-associated protein modules (44, 45). The majority of the previously identified JBTS-associated proteins participate in specific sub-modules of complex ciliary protein networks that vary in sub-ciliary localization, concentrating at the transition zone to organize and regulate the ciliary gate (40, 46).

Using a combination of affinity proteomics (TAP) and Y2H protein interaction screens, we found that JBTS-associated protein ARMC9 forms a complex with the known JBTS-associated proteins CSPP1 and CEP290, not previously known to physically and functionally connect, confirming the importance of this complex to JBTS. We also identified two ciliary microtubule associated proteins that, like CSPP1, directly interact with ARMC9: TOGARAM1 and CCDC66. Although neither of these proteins have previously been implicated in human disease, CCDC66 was found to interact with CEP290 (28) and null mutations cause retinal degeneration in dogs (47) and mice (48). A subsequent TAP experiment using TOGARAM1 as a bait yielded a complex that indeed contained ARMC9, further validating their interaction, but also included another JBTS-associated protein, CEP104 (JBTS25). CEP104 localizes both to the daughter centriole as well as to the apical tip of a growing cilium (22) and, similar to TOGARAM1, interacts with tubulin through its TOG domain (49). Moreover, it interacts with the MT plus end tracking protein MAPRE1 (EB1), with the ciliogenesis regulator CP110 (27), and with NEK1 (49) associated with the ciliopathy “short-rib polydactyly syndrome Majewski type” (50). Co-IP and yeast two-hybrid analysis validated a complex of the core module, ARMC9, TOGARAM1, CCDC66, and CEP104. By Y2H screening we determined the direct interaction of TOGARAM1 with another JBTS-associated protein, RPGRIP1L (JBTS7), while a TAP experiment using CSPP1 as a bait again identified CEP290 as a complex member, and confirmed its previously identified interaction with RPGRIP1L (51). Our results are in agreement with the BioID proximity interactome of CEP104, that contained most our module components, except for ARMC9 and RPGRIP1L (42). Following the “guilt by association” paradigm, we next found biallelic *TOGARAM1* variants in multiple individuals with JBTS, reiterating the relevance of the complex to JBTS, and moving us closer to identifying all genetic causes of this disorder. We did not find *CCDC66* variants in >500 families affected by JBTS, indicating that variants in this gene are, at most, a very rare cause of JBTS.

### Role of the ARMC9-TOGARAM1 complex in ciliary length and stability

The interaction between ARMC9 and TOGARAM1 is contingent on an intact TOGARAM1 TOG2 domain. The structure of TOG domains is highly conserved for microtubule binding, where the intra-HEAT loop within the discontinuous TOG domain binds tubulin (52–55). TOG domains are thought to function to positively regulate microtubule growth and dynamics (56). Interestingly, we found that overexpression of wild-type TOGARAM1 causes increased ciliary length in control and TOGARAM1 mutant RPE1 cells (Summary Figure 9). It has previously been shown that different TOG domains in TOGARAM1 have differential microtubule binding capacity and likely function in concert to coordinate microtubule polymerization (26). The C-terminal TOG domains TOG3 and TOG4 promote microtubule lattice binding (26). In hTERT-RPE1 cells, the TOG2 domain variants Arg368Trp and Leu375Pro interfere with interaction between overexpressed TOGARAM1-and ARMC9; however TOG2 variant TOGARAM1 protein localization and ciliary length remain comparable to wild-type TOGARAM1. TOGARAM1 with TOG3 domain variant Arg1311Cys maintains its interaction with ARMC9 while producing severely shortened cilia, suggesting that TOGARAM1 without a functional TOG3 domain may be unable to support microtubule lattice binding effectively.

Cilia are shorter in fibroblasts from patients with ARMC9- and TOGARAM1-associated JBTS. In contrast, overexpression of wild-type TOGARAM1 causes longer cilia, while overexpression of TOGARAM1 with JBTS-associated variants, can cause longer (TOG2 domain variants) or shorter (TOG3 domain variant) cilia (Summary Figure 9). This is consistent with a dominant negative effect of the TOG3 domain variant, since this effect is observed in control cells expressing endogenous TOGARAM1. Since both long and short cilia have been identified in fibroblasts from patients with different genetic causes of JBTS, no simple correlation between cilium length and JBTS disease mechanism can be made (23, 24, 57, 58). Work in Tetrahymena indicated that TOGARAM1 and ARMC9 orthologs may have opposite effects on B-tubule length (21). In mammalian cells however, we found that mutations in either gene lead to similar shorter cilia, decreased post-translational modifications and increased cold-induced ciliary microtubule depolymerization, suggesting reduced ciliary stability with mutations in either gene. Intriguingly, dysfunction of TOGARAM1 and ARMC9 seem to have opposite effects on the kinetics of cilium resorption after serum re-addition in patient fibroblasts. This result suggests that different mechanisms may underlie ciliary resorption in the setting of serum re-addition versus cold-induced depolymerization. The latter may represent an acute stressor directly correlated with cilium stability, while the former is certainly a regulated mechanism required for cell cycle reentry, for which TOGARAM1 and ARMC9 may indeed play opposing roles as suggested by the work in *Tetrahymena* (21).

### Post-translational modifications of ciliary microtubules

*ARMC9* and *TOGARAM1* dysfunction leads to significantly decreased axonemal post-translational modifications (PTMs, polyglutamylation and acetylation) in patient fibroblasts and zebrafish, supporting the relevance of altered PTMs in JBTS. Tubulin PTMs are indispensable for proper microtubule function, affecting their mechanical properties, stability, and binding of microtubule-associated proteins (MAPs) to influence protein trafficking and signaling (13).

Polyglutamylation decorates the surface of axonemal microtubules. This reversible modification ranges from 1-17 glutamyl residues *in vivo* (59), and plays a role in IFT activity and MAP binding (60–64). Decreased ciliary polyglutamylation interferes with kinesin2-mediated anterograde transport, also on the B-tubule, and subsequently negatively impacts Hedgehog signaling (61, 63). Some MAPs are sensitive to the amount of glutamylation. For example, spastin has optimal microtubule-severing activity in vitro with moderate polyglutamylation, with both hypo-and hyper-glutamylation suppressing activity (65). In the context of JBTS, decreased axonemal polyglutamylation was reported in fibroblasts from patients with *CEP41*-related JBTS (66). More recent work found decreased axonemal glutamylation with *ARL13B*, *FIP5* and *TTLL5* knockdown in immortalized cells, associated with impacts on polycystin localization and Hedgehog signaling (60).

Acetylation of ciliary microtubules is important for motor protein binding and movement, stability, and ciliary disassembly, where deacetylation is required for resorption (67). While most PTMs are added to the C-terminus of tubulin on the microtubular surface, acetylation uniquely occurs on the luminal surface of -tubulin. It has been long observed that the hyperstabilized ciliary microtubules are acetylated, but until recently it was not known if the modification confers stability or if long-lived stable microtubules accumulate this modification. Recent work using cryo-EM confirmed that acetylation causes a stabilizing conformational change (68). This is in line with our findings of decreased axonemal microtubule acetylation and stability with ARMC9 and TOGARAM1 dysfunction, as well as previously published findings with Kif7 dysfunction (57). The more rapid ciliary resorption with ARMC9 dysfunction is unlikely due to excessive deacetylation since cilia were not stabilized by HDAC6 inhibition. Interestingly, Kif7 mutant mouse embryonic fibroblasts (MEFs) exhibited reduced glutamylation, making it the only other JBTS model with decreases in both PTMs (23, 57). Fibroblasts from patients with *INPP5E*-related JBTS also display decreased ciliary stability (69). Notably, these models of KIF7 and INPP5E ciliary dysfunction disrupt Hh signaling, likely due to GLI/SUFU mislocalization and aberrant phosphatidylinositol composition respectively, but it is unknown if/how PTMs also contribute to Hh dysfunction.

The observed reduction of PTMs with ARMC9 and TOGARAM1 dysfunction could therefore affect ciliary function through loss of stability and/or direct disruption of signaling pathways. In fact, *Armc9* was identified as a positive regulator of the Hh pathway in a genome-wide screen for Hh signaling components (25). That study also demonstrated that over-expressed ARMC9 translocates from the ciliary base to the distal tip upon Hh pathway stimulation (25). In *Tetrahymena*, orthologs of ARMC9 and TOGARAM1 are seen at both the base and tip, with tip enrichment during cilia regeneration (21). Taken together, these results suggest a dynamic localization of the complex members, and likely changes in protein complex composition at each locale. Further work will be required to determine the dynamic localization of the ARMC9-TOGARAM1 complex during ciliogenesis, resorption, and signaling.

## Conclusions

The biological mechanisms underlying JBTS remain incompletely understood. This work brings us one step closer to the complete catalog of JBTS genetic causes, and highlights the role of a new JBTS-associated protein complex including ARMC9 and TOGARAM1. Approximately half of JBTS-associated genes are implicated in transition zone function which is required for ciliary ARL13B and INPP5E localization. In contrast, the ARMC9-TOGARAM1 complex is not required for INPP5E localization, and instead, appears to regulate the post-translational modification of ciliary microtubules, ciliary length and ciliary stability. Future work will need to reconcile how the diverse array of cellular defects associated with loss of function for the JBTS genes relate to this important human disorder.

## Materials and Methods

### Patient Fibroblasts Culture

Patient fibroblasts were cultured at 37⁰C, 5% CO_2_ in DMEM (Gibco, 11995-065) supplemented with 10% FBS, 1% penicillin/streptomycin. Fibroblasts are seeded at 5×10^5 and allowed to reach 70-80% confluency prior to passages. For experiments, cells are serum starved in DMEM only for 48 hours to induce ciliogenesis. All cell lines used were within 1 passage number of each other and ≤10 total passages. All cell lines were routinely tested for mycoplasma.

### CRISPR/Cas9 generation of *TOGARAM1* mut hTERT-RPE1 lines

hTERT-RPE1 were maintained in culture according to ATCC specifications. Cells were plated in 1 well of a 6 well plate. At ∼70-80% confluency, cells were transfected with the Cas9 backbone px459v2 containing gRNAs to TOGARAM1 using lipofectamine 2000 (Thermo Fisher). Sequencing primers for genotyping targeted the 5’UTR CTGAAGCTGTTCTTTTGCCTCT (forward) and exon 1 CTACCTCCTTCCACAAGCACTC (reverse). Both TOGARAM1 mut clone 1 and 2 have compound heterozygous mutations which result in large deletions including the ATG site and a portion of exon 1 (Supplemental Figure 4 A, B). Cloning was performed as previously described (70). gRNAs to the 5’UTR and ATG site (5’- CACCTGACAACCCTGCATGG-3’) and exon 1 (5’-TCTGGAGGCGGTTTGTCAGG-3’), were designed utilizing CHOPCHOP. pSpCas9(BB)-2A-Puro(PX459)V2.0 from Feng Zhang (Addgene plasmid # 62988; http://n2t.net/addgene:62988; RRID:Addgene_62988) was purchased from Addgene.

### hTERT-RPE1 Immunofluorescence

hTERT-RPE1 cells were cultured according to ATCC specifications. For immunofluorescence imaging, cells were plated on glass coverslips. At 24hr after plating, cells were serum-starved for 48hr in 0.2% FBS medium, transfections were performed where indicated at 48 hours post plating using lipofectamine 2000 (Thermo Fisher) in accordance with the manufacturer’s protocol. 72 hours after plating, cells were rinsed once with 1X PBS, fixed with 2% paraformaldehyde for 20 min and permeabilized with 1% Triton-X for 5 min. All steps were performed at room temperature. Cells were blocked in 2% BSA for 45 min, then incubated with the following antibodies for 1hr: anti-ARMC9 (rabbit, Atlas Antibodies cat# HPA019041, RRID: AB_1233489; 1:200), anti-ARL13B (rabbit Proteintech, cat. no. 17711-1-AP; 1:500), anti-RPGRIP1L (guinea pig, in house; SNC040, 1:300), anti-acetylated tubulin antibody (mouse, clone 6-11-B1, Sigma-Aldrich, cat. no. T6793; 1:1,000), and anti-GT335 (mouse, a kind gift from Carsten Janke; 1:2000). Cells were stained with secondary antibodies for 45 min. The secondary antibodies used were (all from Life Technologies/Thermo Fisher Scientific; diluted 1:500 in 2% BSA): anti-guinea pig IgG Alexa Fluor 647, anti-rabbit IgG Alexa Fluor 488, and anti-mouse IgG Alexa Fluor 568. Fluoromount-G with DAPI (ThermoFisher) was used to mount the cover slips. Imaging was performed with the Zeiss Axio Imager Z2 Microscope or the Zeiss LSM 880 Laser scanning confocal microscope equipped with Airyscan technology.

### Patient fibroblast immunofluorescence

Patient fibroblasts were grown on coverslips (Neuvitro, GG-12-1.5.pdl, 0.3mg/mL Poly-D-lysine coating, 1.5mm thickness), serum-starved for 48 hours, then fixed in ice cold 2% paraformaldehyde for 20 minutes. After a PBS wash, cells were permeabilized with 1% Trition-X in PBS or ice-cold Methanol for 5 minutes. Cells were blocked in 2% BSA in PBS for 1 hour at room temperature, then incubated with the following antibodies (in 2% BSA/PBS) for 1.5 hours at room temperature: mouse anti-glutamylated tubulin, GT335, (Adipogen, AG- 20B-0020-C100, 1:2000), mouse anti-acetylated tubulin (clone 6-11-B1, Sigma-Aldrich, T6793; 1:1,000), mouse anti-ARL13B (UC Davis NeuroMab 75-287 clone N295B/66, 1:2000), goat anti-γ- tubulin (Santa Cruz, SC-7396 1:200), rabbit anti-ARL13B (Proteintech, 17711-1-AP, 1:200), or rabbit anti-INPP5E (Proteintech, 17797-1-AP, 1:100). Cells were washed thrice with PBS and stained with secondary antibodies (in 2% BSA/PBS) for 1 hour at room temperature (all Invitrogen at 1:2000, anti-goat-647, A21447, anti-rabbit-488, A11008). After three PBS washes, coverslips were mounted on slides using Fluoromount-G with DAPI (Invitrogen, 00-4959-52) and sealed with nail polish.

### Patient Fibroblasts Microscopy & Immunofluorescence Quantification

Wide-field fluorescent images were acquired on an 3i imaging workstation (3i, Denver, CO) with Axio inverted microscope with Definite Focus (Zeiss). For each experiment, optimal exposures were determined for each fluorophore to ensure that we used the full dynamic range of our CoolSnap HQ2 camera (Photometrics, Inc., Tuscon, AZ) without saturating any pixels. Dark field correction was applied to each channel to remove artifacts generated from the camera and electronics due to non-uniformities in illumination. Z-stack images with 0.3 μm steps were acquired at ≥10 distinct locations on each slide with a 40x objection using identical scope settings for all slides in an experiment. Sum projected images were analyzed in FIJI (NIH). A reference ciliary mask was drawn atop the reference signal for each cilium by standardized methods. A skeleton measurement of this mask extracted ciliary length data. The average fluorescence intensity was measured within the cilium mask in the channel of interest. To correct for antibody background, the background from a region directly adjacent to each cilium was measured and subtracted.

### Zebrafish: Phylogeny & Synteny Analysis, CRISPR gene editing, Scanning electron microscopy & Immunofluorescence

Phylogeny and Synteny analyses were performed as previously described (71), using the Phylogeny.fr platform (http://www.phylogeny.fr/) and the synteny database (http://syntenydb.uoregon.edu/synteny_db/). Briefly, for Phylogeny, length of input sequences varied between 256 (*xenopus* truncated version) and 516 amino acids, and after curation 487 amino acids were used for further analysis. For Synteny analysis, parameters were adjusted to sliding window sizes between 25 and 100, and several genes in the vicinity of *Togarams* were used for additional syntenic comparison. Zebrafish (*Danio rerio*) were maintained at 28 °C with a 14 h/10 h light/dark cycle as previously described (ref. M. Westerfield 1993). All zebrafish protocols were in compliance with internationally recognized guidelines for the use of zebrafish in biomedical research, and the experiments were approved by local authorities (Veterinäramt Zürich TV150). Generation and genotyping of the *armc9* mutant zebrafish was previously described (24). sgRNAs for *togaram1* CRISPR/Cas9 mutagenesis were designed with the website CHOPCHOP: 3’- GGGGTCTCCTCTGCTGGGCC-5’ and 5’-GGACGAGATGCTGGACCGAG-3’ for exon 5/6 and 3’- GGCTGCCGATGACCAGAGCT-5’ and 5’-GGTGAATCTGCGCGCTCTGG-3’ for exon 20/21 in *togaram1.* sgRNAs were mixed with Cas9 protein (gift from Darren Gilmour, M0646M NEB, or B25641 invitrogen) and co-injected into 1-cell stage embryos using a microinjector (Eppendorf). Amplification of the target regions for genotyping was performed using primer pairs 5’-AGACGCTCCTCAACTCCAGA-3’ and 5’- GCCGTGTAGACGAGTGTGTT-3’ for exon 20/21 in *togaram1.* The PCR products were analyzed with gel electrophoresis and subcloned before sequencing. Experiments were performed using the *armc9^zh505^* and the *togaram1^zh509^* or *togaram1^zh510^* mutant (Supplemental Figure 5F). Zebrafish larvae were fixed in 2.5 % Glutaraldehyde in 0.1 M Cacodylate buffer and prepared for scanning electron microscopy (SEM) following standard protocols. SEM was performed on a ZEISS Supra VP 50 microscope. Whole-mount immunohistochemistry was performed on zebrafish larvae fixed in 4% paraformaldehyde or 80% MeOH in DMSO according to standard protocols. The following primary antibodies were used: acetylated tubulin (1:400, Sigma 7451), GT335 (1:400, Enzo Life Sciences A20631002), arl13b (1:100, gift from Z.Sun (72)), cc2d2a (73). Images were taken with a Leica HCS LSI confocal microscope. Acetylated tubulin and glutamylated tubulin mean fluorescence intensity was quantified using FIJI: fluorescence intensity of 10 cilia from each larvae were measured and averaged, so that each datapoint in graphs Fig 6A, 7E and 7F represents one individual larva. The background was subtracted from each measurement.

### Cloning

All expression constructs were generated using Gateway Technology (Life Technologies) according to manufacturer’s instructions. The constructs generated encoded TOGARAM1 (and JBTS associated variants), ARMC9, CCDC66, and CSPP1 in the following destination vectors: 3xHA, 3xFlag, TAP, myc, mRFP, GAL4-BD, and PalmMyr-CFP. The entry clone of human TOGARAM1 (<u>NM_001308120.2</u>) was synthesized and purchased from VectorBuilder. Constructs encoding TOGARAM1 and variants were generated by site directed mutagenesis PCR. All entry clone sequences were verified using Sanger sequencing.

### PalmMyr Assay

hTERT-RPE1 cells were plated on glass slides, 24 hours later when cells reached approximately 80% confluency they were transfected using lipofectamine 2000 (ThermoFisher) with either mRFP-tagged TOGARAM1 / TOGARAM1 variants or PalmMyr-CFP-tagged ARMC9 or both. 24 hours post transfection, cells were starved for an additional 24 hours, fixed with 2% PFA at room temperature and prepared for analysis. Imaging was performed with the Zeiss Axio Imager Z2 Microscope.

### Tandem affinity purification & mass spectrometry

HEK293Tcells were maintained according to ATCC specifications. Cells were seeded and expanded for 16 – 24 hours, then transfected with the corresponding SF-TAP-tagged DNA constructs using PEI reagent (Polysciences) according to manufacturer’s protocol. 48 hours later, cells were harvested in lysis buffer containing 0.5% Nonidet-P40 (NP-40), protease inhibitor cocktail (Roche), and phosphatase inhibitor cocktails II and III (Sigma-Aldrich) in TBS (30mM Tris-HCl, pH 7.4 and 150mM NaCl) for 20 min at 4°C. Cell debris and nuclei were removed by centrifugation at 10,000g for 10 min. For SF-TAP analysis, the cleared supernatant was incubated for 1 hour at 4°C with Strep-Tactin superflow (IBA). Subsequently, the resin was washed three times in wash buffer (TBS containing 0.1% NP-40 and phosphatase inhibitor cocktails II and III, Sigma-Aldrich). Protein baits were eluted with Strep-elution buffer (2mM desthiobiotin in TBS). For the second purification step, the eluates were transferred to anti-Flag M2 agarose beads (Sigma-Aldrich) and incubated for 1 hour at 4°C. The beads were washed 3x with wash buffer and proteins were eluted with FLAG peptide (200 mg/ml, Sigma-Aldrich) in TBS. After purification, the protein samples were precipitated with chloroform and methanol and subjected to in-solution tryptic cleavage. Mass spectrometry and analysis were performed as previously described (43).

### Yeast two-hybrid

The GAL4-based yeast two-hybrid system was used to screen for binary protein–protein interactions. Yeast two-hybrid constructs containing the cDNAs of selected bait proteins ARMC9, TOGARAM1, CCDC66 and fragments thereof were generated by Gateway adapted PCR and cloning. Constructs encoding full-length or fragments of bait proteins fused to a DNA binding domain (GAL4-BD) were used to screen human oligo-dT primed and bovine random hexamer primed retinal cDNA libraries, prey proteins are fused to a GAL4 activation domain (GAL4-AD). The yeast strain PJ96-4A, which carries the HIS3 (histidine), ADE2 (adenine), MEL1 (a-galactosidase) and LacZ (b-galactosidase) reporter genes, was used as a host. Interactions were analyzed by assessment of reporter gene (HIS3 and ADE2) activation via growth on selective media and β-galactosidase colorimetric filter lift assays (*LacZ* reporter gene). cDNA inserts of clones containing putative interaction partners were confirmed by Sanger sequencing.

### Coimmunoprecipitation

HEK293Ts were plated in 6 well plates and transfected using Effectene Transfection Reagent (Qiagen) according the manufacturer’s protocol. 24 hours post-transfection cells were lysed in 200 uL/well of ice cold IP lysis buffer and collected for centrifugation. Lysates were then incubated in Eppendorf tubes with HA affinity matrix beads (Roche). Lysates were nutated for 2 hours at 4°C. Beads were washed 3x in 1 ml of ice cold IP lysis buffer, all liquid was removed from the beads using a syringe with 0.5 mm needle. 50 uL of NuPAGE loading dye plus 100 mM DTT was added to the samples and they were heated at 95⁰C for 10 minutes. For the co-IP performed in the ARMC9 IMCD3 FlpIn lines (25) as shown in figure 1c, cells constitutively expressing mouse ARMC9-3xFlag were transfected with HA-TOGARAM and subsequently co-immunoprecipitated with HA beads as described above. Western blotting was performed using the standard protocol for the NuPAGE system and visualized on the Odyssey. c-myc (Roche, 11667149001; 1:500), HA (Sigma; H9658; 1:1000), and Flag (Sigma; F3165-0.2MG; 1:1000) primary antibodies were used. The secondary antibody used was goat anti-Mouse IRDye800 (Licor biosciences; 926-32210; 1:20,000).

### Western blotting

Trypsinized and concentrated cells were lysed with NP-40 cell lysis buffer (ThermoFisher, FNN0021). Cellular proteins were denatured using Laemmli sample buffer supplemented with 2-Mercaptoethanol (BioRad, #1610747, #1610710) and heated at 95⁰C for 10 minutes. Cellular proteins were separated by SDS-PAGE and transferred to a PVDF membrane (Millipore IPVH00010) using standard protocols. The following primary antibodies were used: mouse anti-acetylated tubulin (clone 6-11-B1, Sigma-Aldrich, T6793; 1:1,000), mouse-anti beta-actin (Sigma, A5441, 1:5000), rabbit anti-ARL13B (Proteintech, 17711- 1-AP, 1:1000), rabbit anti-INPP5E (Proteintech, 17797-1-AP, 1:1000), rabbit anti-Giantin (Abcam, ab24586, 1:5000) or rabbit anti-ARMC9 (N-terminal epitope, Atlas Antibodies cat# HPA019041, RRID: AB_1233489; 1:2000). Western blots were developed using anti-mouse or anti-rabbit secondary antibodies conjugated to horseradish peroxidase (anti-rabbit-HRP, Novex A16029 1:2000, anti-mouse-HRP, Invitrogen G21040 1:2000) and chemiluminescent substrate (BioRad Clarity Max 1705062). A ChemiDoc MP imaging system with ImageLab software was used for imaging (BioRad).

### Statistics & reproducibility

Statistical analyses were performed in Excel and Graphpad/Prism6. Graphical data presented as percentages include 95% confidence intervals. Quantitative immunofluorescence statistics were calculated with Student’s t-test with unequal variances. Fluorescence intensity measurements from multiple experiments were combined for statistical analysis. Experiments were independently performed thrice. In cilia stability assays, linear regression of ciliary loss over time yielded the slope, and we used students T-test with unequal variances for singular comparisons and a one-way ANOVA to assess significance for multiple samples. N values are stated in the Figures and either represent cilia or cells (as indicated in each Figure). P values are stated in the figure legends, and symbols indicate the following P values: ns, P > 0.05; *, P ≤ 0.05; **, P ≤ 0.01; ***, P ≤ 0.001; ****, P ≤ 0.0001; *****, P ≤ 0.00001.

## Study approval

### Microtubule cold assay

For microtubule cold depolymerization assays, cells were grown until near-confluent on coverslips in 24- well plates, serum starved for 48 hours, then placed at either room temperature or 4⁰C for 10 minutes. Cells were then fixed as described and processed for immunofluorescence to determine ciliation percentage.

### Stability assay

For cilia stability assays, cells were grown until near-confluent on coverslips in 24-well plates, serum starved for 48 hours, then media was replaced with DMEM 10% FBS at hourly time points for 4-8 hours. Cells were then fixed as described and processed for immunofluorescence to determine ciliation percentage. In a subset of these experiments, we blocked histone deacteylase 6 (HDAC6) activity, with a specific inhibitor, tubacin (Sigma, SML0065). Cells were either treated with 1 μM tubacin in DMSO or in DMSO only as a vehicle control.

### Subject Ascertainment & Phenotypic Data

Participants were enrolled under approved human subjects research protocols at the University of Washington (UW) or King Faisal Specialist Hospital and Research Centre KFSHRC RAC# 2070023, 2080006 and 2121053. All participants or their legal guardians provided written informed consent. All participants have clinical findings of JS (intellectual impairment, hypotonia, ataxia, and/or oculomotor apraxia) and diagnostic or supportive brain imaging findings (MTS or cerebellar vermis hypoplasia). Clinical data were obtained by direct examination of participants, review of medical records, and structured questionnaires.

### Variant Identification

Samples from individuals affected by JS were previously screened using a molecular inversion probes (MIPs) targeted capture (74). For gene list see supplemental table 4 (6, 7, 9, 14, 15, 20, 23, 32, 40, 45, 75- 89). In samples without causal variants, exome sequencing was performed as previously described (90) using Roche Nimblegen SeqCap EZ Human Exome Library v2.0 capture probes (36.5 Mb of coding exons) and paired-end 50 base pair reads on an Illumina HiSeq sequencer. In accordance with the Genome Analysis ToolKit’s (GATK) best practices, we mapped sequence reads to the human reference genome (hg19) using the Burrows-Wheeler Aligner (v.0.7.8), removed duplicate reads (PicardMarkDuplicates v.1.113), and performed indel realignment (GATK IndelRealigner v.3.1) and base-quality recalibration (GATK TableRecalibration v.3.1). We called variants using the GATK UnifiedGenotyper and flagged with VariantFiltration to mark potential false positives that did not pass the following filters: Heterozygous Allele Balance (ABHet) > 0.75, Quality by Depth > 5.0, Quality (QUAL) > 50.0, Homopolymer Run (Hrun) < 4.0, and low depth (< 8x). We used SeattleSeq for variant annotation and the Combined Annotation Dependent Depletion (CADD) score to determine deleteriousness of identified missense variants (91). Based on CADD score data for causal variants in other JS-associated genes, we used a CADD score cutoff of 15 to define deleterious variants (32)

For the one family in the Saudi Arabian cohort, DNA from the affected individuals, unaffected siblings, and parents were genotyped using the Axiom SNP Chip platform to determine the candidate autozygome (92, 93). WES was performed using TruSeq Exome Enrichment kit (Illumina) following the manufacturer’s protocol. Samples were prepared as an Illumina sequencing library, and then the sequencing libraries were enriched for the desired target using the Illumina Exome Enrichment protocol. The captured libraries were sequenced using an Illumina HiSeq 2000 Sequencer. The reads were mapped against UCSC hg19 by BWA. SNPs and indels were detected by SAMTOOLS. Homozygous rare, predicted-deleterious, and coding/splicing variants within the autozygome of the affected individual were considered as likely causal. We defined rare variants as those with frequency of <0.1% in publicly available variant databases (1000 Genomes, Exome Variant Server, and gnomAD) as well as a database of 2,379 in-house ethnically matched exomes, and defined deleterious if predicted to be pathogenic by PolyPhen, SIFT, and CADD (score > 15).

### Array CGH

To assess copy-number variation, we performed array comparative genomic hybridization using a custom 8×60K oligonucleotide array (Agilent Technologies)(94). For gene list see supplemental table 5. Probe spacing was a median of 11 bp in the exons, and a median of 315 bp throughout the intronic regions and 100 kb on either side of each gene. Data were generated on an Agilent Technologies DNA Microarray Scanner with Surescan High-Resolution Technology using Agilent Scan Control software and were processed and analyzed using Agilent Feature Extraction and Agilent Cytogenomics software. To determine the effect of the deletion in UW360-3, we extracted RNA (BioRad, Aurum Total RNA kit, 7326820) from the associated cell line and converted it to cDNA (Biorad, iScript Reverse Transcription Supermix, 1708840) for downstream Sanger sequencing.

## Web resources

http://chopchop.cbu.uib.no

http://gnomad.broadinstitute.org

https://www.ensembl.org/index.html

http://www.cmbi.umcn.nl/hope/

https://huygens.science.uva.nl/PlotsOfData/(95)

http://www.phylogeny.fr/

http://syntenydb.uoregon.edu/synteny_db/

## Acknowledgements

We thank the patient and parents for their participation in this research. We thank Dr David Breslow (Yale University) for providing the ARMC9-3xFlag IMCD3 FlpIn lines. The research leading to these results has received funding from the Netherlands Organisation for Scientific Research (NWO Vici-865.12.005 to R.R.), the Netherlands Organisation for Health Research and Development (ZonMW, #91216051 to R.R.), the Foundation Fighting Blindness (PPA-0717-0719-RAD to M.U. and R.R.), the Tistou & Charlotte Kerstan Stiftung (to M.U.), the Swiss National Science Foundation SNF (PP00P3_170681 to R.B.G and 31003A_173083 to S.C.F.N.), the Zurich Neuroscience Center (ZNZ) to R.B.G., the NIH Eunice Kennedy Shriver National Institute of Child Health and Human Development (U54HD083091 Genetics Core and sub-project 6849 to D.D. and F32 HD095599 to J.C.V)), private donations (to D.D.), NIH National Human Genome Research Institute (U54HG006493 to M.B. and D.N.) and King Salman Center for Disability Research (FSA).

## Author contributions

R.B.-G., R.R. and D.D. conceived the overall project. B.L., J.V.D.W., T.D.S.R., J.C.D., S.C.F.N., K.B., M.U., F.S.A., R.B.-G., R.R., & D.D. designed experiments and led the data generation and processing. B.L., J.V.D.W., T.D.S.R., A.G., S.J.F.L., M.E.G., S.E.C.v.B., C.M., and U.W.C.M.G. performed experiments. R.S., H.M., & J.C.D acquired clinical phenotype data. B.L., J.V.D.W., T.D.S.R., M.G., M.E.G., C.M., R.B.-G., R.R., & D.D. analyzed and interpreted data. B.L., J.V.D.W., T.D.S.R., R.B.-G., R.R., & D.D. wrote the paper with input from all authors.

