## supplemental figures and legends for "ARMC9 and TOGARAM1 define a Joubert syndrome-associated protein module that regulates axonemal post-translational modifications and cilium stability"

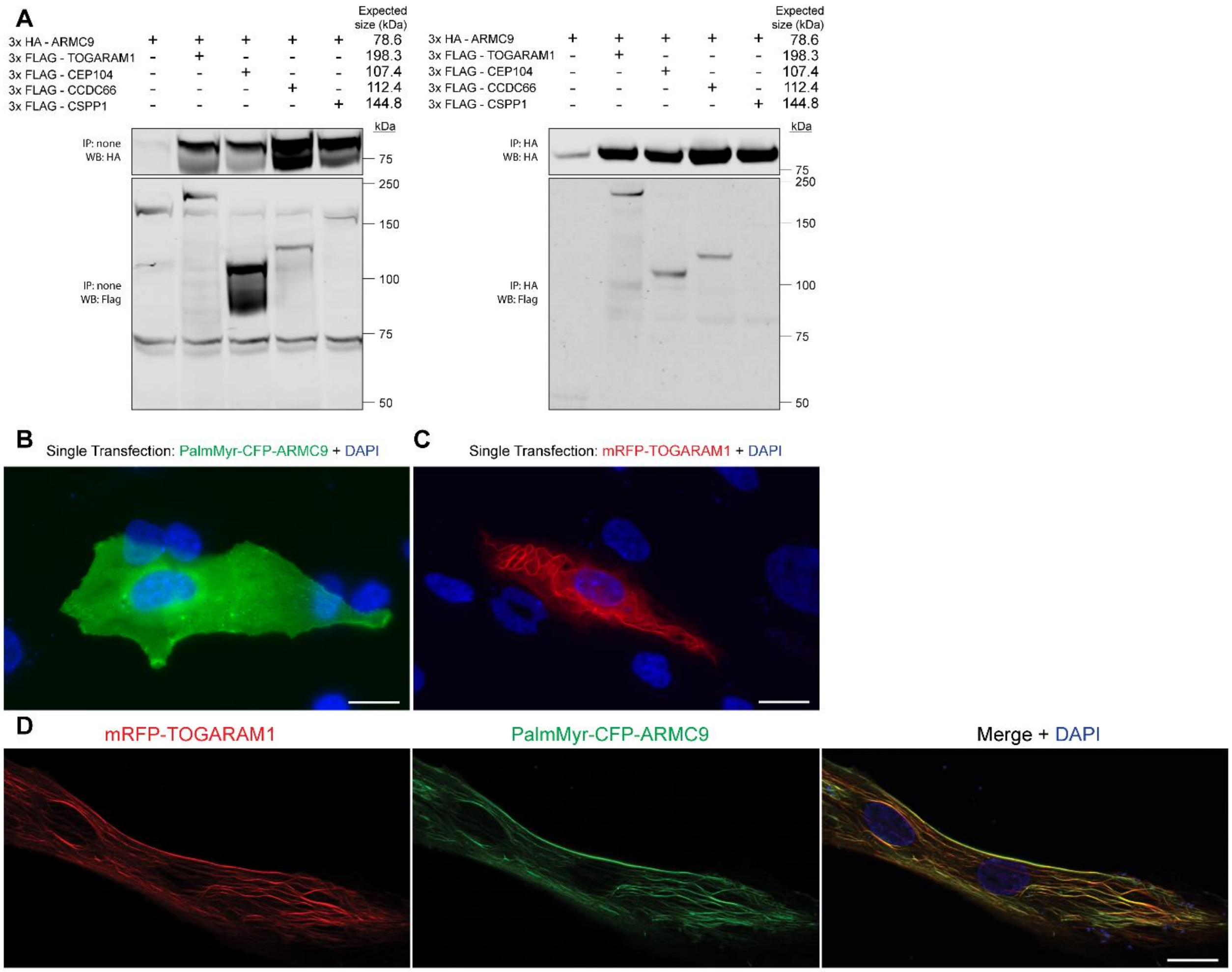


**Supplemental Figure 1** Validation of ARMC9 interactome. **(A)** Co-IP of HA tagged ARMC9 with Flag tagged interactors. HA tagged ARMC9 was transfected alone or in conjunction with Flag tagged interactor constructs, HA beads were used to pull down the bait construct and blots were probed for the presence of the Flag tagged interactors. Experiment was performed in biological triplicates. Full blots are viewable in Plemental Figure 10. **(B)** Single transfection of PalmMyr-CFP-ARMC9 (green) and **(C)** single transfection of mRFP-TOGARAM1 (red) shows the localization in the absence of the respective interactor. **(D)** Co-expression of mRFP-TOGARAM1 and PalmMyr-CFP-ARMC9 shows microtubule colocalization. Scale bar indicates 20 μm.

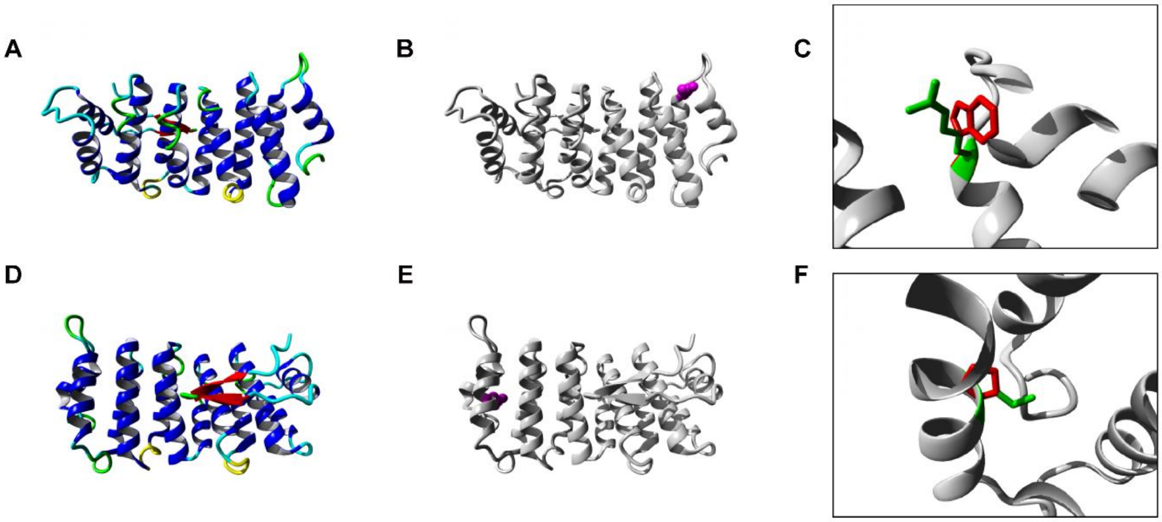


**Supplemental Figure 2** In silico modeling of the TOG2 variants. **(A)** Ribbon model of the wild type TOG2 domain structure of TOGARAM1 generated using HOPE. The protein is colored by the following elements: alpha-helices are show in blue, beta-strand is shown in red, turns are green, 3/10 helix is yellow, and random coil is cyan. **(B)** p. Arg368Trp missense variant in the TOG2 domain in ribbon-presentation generated using HOPE. TOG2 is shown in grey, the side chain of the mutated residue is shown in magenta. **(C)** A close up of the side chain of both wild type and mutated residue is shown in green and red respectively. **(D)** Inverted view of wild type TOG2 domain structure of TOGARAM1 as compared to (a). **(E)** p.Leu375Pro missense variant modeled in the TOG2 domain of TOGARAM1 in ribbon-presentation. TOG2 is shown in grey, the side chain of the mutated residue is shown in magenta. **(F)** A close up of the side chain of both wild type and mutated residue is shown in green and red respectively.

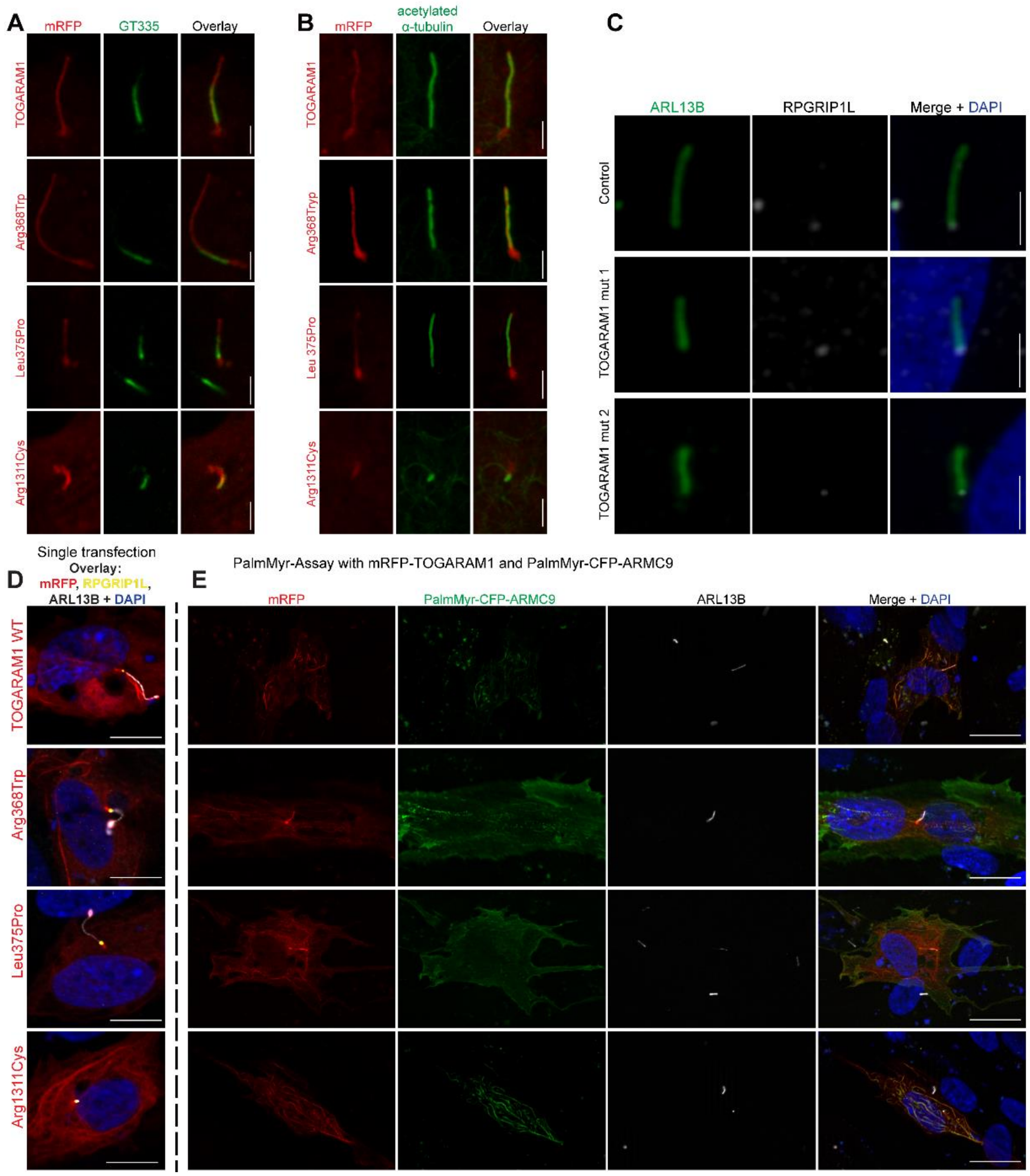


**Supplemental Figure 3** Expression of JBTS-associated TOGARAM1 variants affect ciliary length and the interaction with ARMC9. **(A)** Polyglutamylated **(B)** and acetylated tubulin in cells transfected with mRFP-tagged TOGARAM1 or variants. GT335, shown in green, marks the glutamylated portion of the ciliary axoneme. Acetylated alpha-tubulin, shown in green, co-stains the cilium. All images are representative of 3 independent imaging experiments. Compressed z stacks of polyglutamylated and acetylated alpha tubulin indicate partial or full colocalization with mRFP-TOGARAM1. Scale bars are 2 µm. **(C)** Images of cilia from WT and TOGARAM1 mut RPE1 lines 1 and 2, ARL13B (green), RPGRIP1L is shown in white. Scale bar represent 2 µm. **(D)** PalmMyr assay with ARMC9 and TOGARAM1 in WT hTERT-RPE1 cells: Single transfections of mRFP tagged TOGARAM1 wildtype and variants show localization in the absence of the ARMC9 PalmMyr construct. In the overlay mRFP (red) is shown with ARL13B (white), RPGRIP1L (yellow), and DAPI (blue). Scale bar represents 10 µm. **(E)** Co-expression of mRFP-TOGARAM1 and PalmMyr-CFP-ARMC9: Wild-type and Arg1311Cys TOGARAM1 show colocalization with PalmMyr-CFP-ARMC9, indicating an interaction, while TOGARAM1 variants Arg368Trp and Leu375Pro indicate a highly reduce or absence of interaction. Scale bar indicates 20 µm.

A

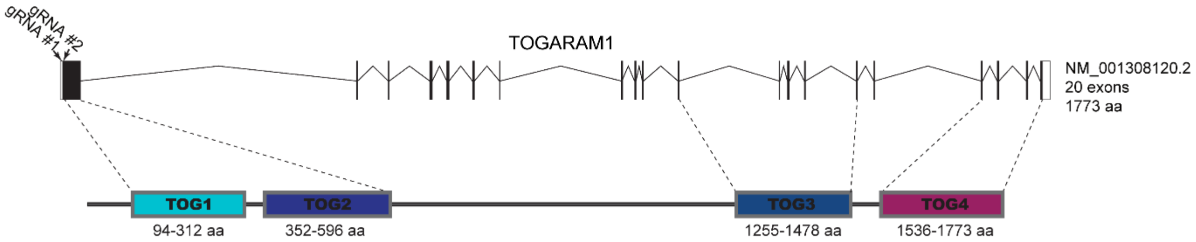


B

hTERT-RPE1 mut lines

| NM_001308120.2 | allele 1 | allele 2 |
|---|---|---|
| TOGARAM1 mut 1 | c.[2_269del;270dup] | c.[2_269inv] |
| TOGARAM1 mut 2 | c.[-33_314del] | c.[2_269inv; 269_270insTC] |

**Supplemental Figure 4** CRISPR/Cas9 edited *TOGARAM1* hTERT-RPE1 mutant lines. **(A)** Schematic representation of the TOG array aligned with the genomic region encoding TOGARAM1. The target sites of gRNA 1 and gRNA 2 are indicated by arrows in exon 1. They are predicted to direct cleavage at cDNA position 2 and 269 respectively. This portion of exon 1 encodes the region of the protein immediately before the TOG1 domain. **(B)** *TOGARAM1* mut 1 harbors a 267 bp deletion and a single bp duplication in one allele and a 267 bp inversion in the other allele, both occurring in exon 1: NM_015091.2:c.[2_269del;270dup]; [2_269inv]. *TOGARAM1* mut line 2 harbors a 347 bp deletion in one allele and a 267 bp inversion and 2 bp insertion in the other allele, both occurring in exon 1: NM_015091.2:c.[-33_314del]; [2_269inv; 269_270insTC].

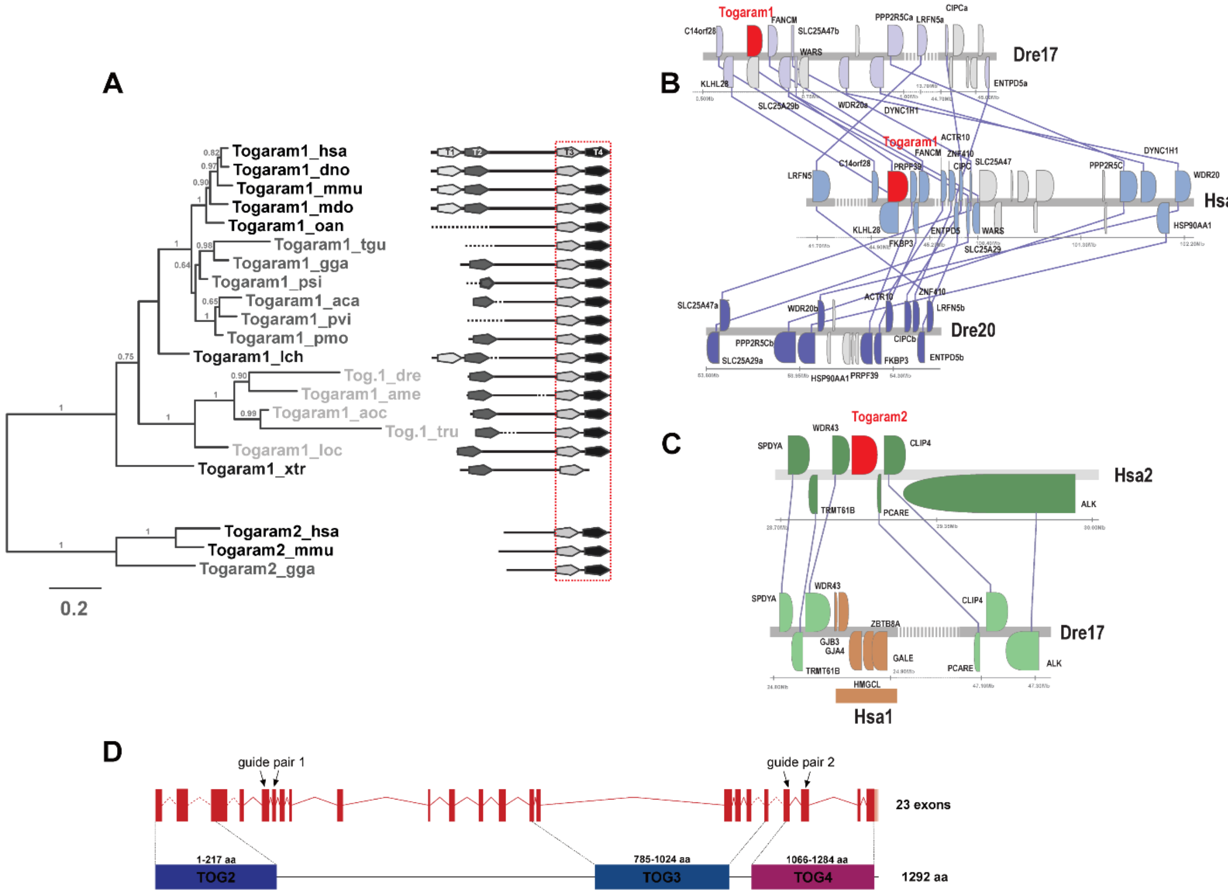


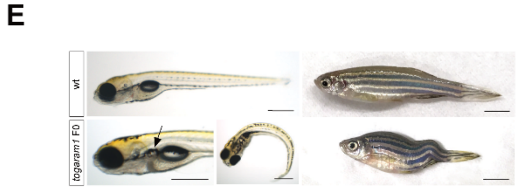


**F**

| allele | mutation, location |
|---|---|
| *armc9*$^{zh501}$ | +8 bp, exon 4 |
| *armc9*$^{zh505}$ | -37 bp, exon4 |
| *armc9*$^{zh503}$ | -10 bp, exon 14 |
| *armc9*$^{zh504}$ | +57 bp, exon 14 |
| *armc9*$^{zh505}$ | +110 bp, exon 14 |
| *togaram1 -1* | -1 bp, exon 20 |
| *togaram1 -31* | -31 bp, exon 20/21 |
| *togaram1*$^{zh508}$ | -21 bp, exon 21 |
| *togaram1*$^{zh509}$ | -37 bp, exon 20/21 |
| *togaram1*$^{zh510}$ | -407 bp, exon 20/21 |

**Supplemental Figure 5** Togaram1 phylogeny, synteny and zebrafish F0 phenotypes. **(A)** Phylogenetic analysis of the C-terminal part (red box) of TOGARAMs in different vertebrate species revealed a clear phylogenetic separation of *TOGARAM1* and *TOGARAM2*. The following species were used for phylogeny: *Anolis carolinensis* (aca), *Astyanax mexicanus* (ame), *Amphiprion ocellaris* (aoc), *Dasypus novemcinctus* (dno), *Danio rerio* (dre), *Gallus gallus* (gga), *Homo sapiens* (hsa), *Latimeria chalumnae* (lch), *Lepisosteus oculatus* (loc), *Monodelphis domestica* (mdo), *Mus musculus* (mmu), *Ornithorhynchus anatinus* (oan), *Pelodiscus sinensis* (psi), *Pogona vitticeps* (pvi), *Taeniopygia guttata* (tgu), *Takifugu rubripes* (tru), *Xenopus tropicalis* (xtr). Dotted lines in the domain representation represent sequence strings so far not covered in the corresponding genome assemblies. In the phylogenetic tree mammals are given in black, amphibians, birds, turtles and reptiles in dark gray and teleosts in light gray. The scale bars represent the distance were 20% of the amino acids are changed. **(B)** Synteny analysis confirms orthology between human and zebrafish *TOGARAM1*. The human *TOGARAM1* gene is located on chromosome 14. Corresponding chromosomal regions to the human chromosome 14 are located on zebrafish chromosomes 17 and 20. In contrast to the zebrafish chromosome 17 where a *TOGARAM1* gene can be readily identified, zebrafish chromosome 20 lacks a corresponding ortholog, suggesting that in the case of *TOGARAM1* no zebrafish duplicate of this gene has been retained. **(C)** Synteny of human *TOGARAM2* shows no corresponding gene in zebrafish. Interestingly, no ortholog of human *TOGARAM2* (located on human chromosome 2) is present in the zebrafish genomic region where the genes flanking *TOGARAM2* are located. **(D)** Zebrafish *togaram1* exons and corresponding protein with domains. Red dashed lines represent unknown intron size. C-terminal transparent part represents unknown stop codon. Location of sgRNAs for genome editing are indicated: two different sgRNAs per target region were co-injected to generate larger deletions. **(E)** Phenotype of the *togaram1* F0 mosaic zebrafish: Larvae have kidney cysts (arrow) and body curvature. Adults develop scoliosis. Scale bars are 500 µm for larvae and 5 mm for adults. **(F)** Alleles generated with CRISPR/Cas9 for *armc9* and *togaram1*.

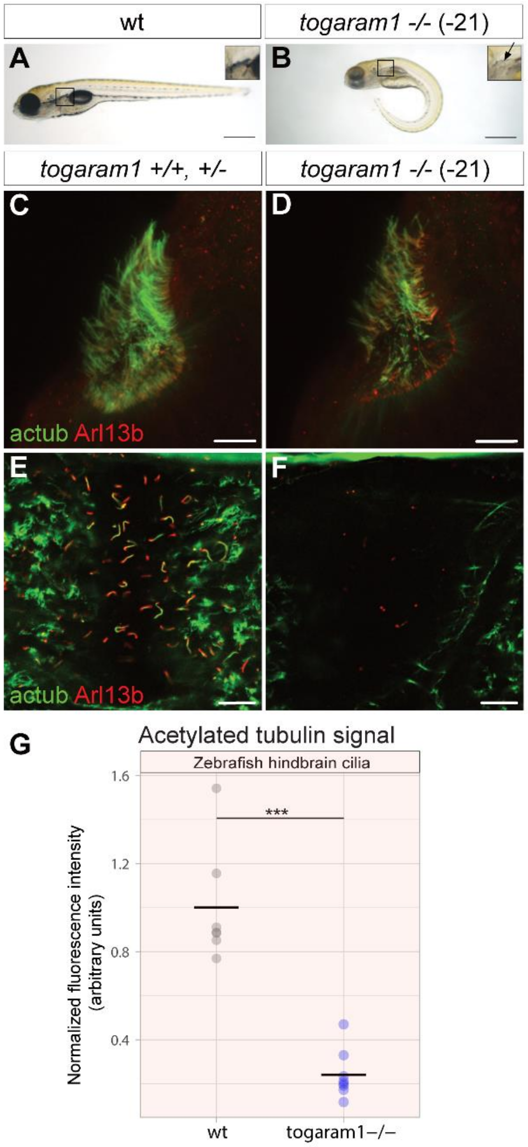


**Supplemental Figure 6** Phenotypes of *togaram1*$^{zh508}$ in-frame mutant zebrafish. **(A)** Wild-type **(B)** and mutant larval phenotype with curved body shape and kidney cysts in *togaram1*-/- (-21). Scale bars are 500 µm. **(C)** Wild type **(D)** and *togaram1*-/- (-21) immunofluorescence of 3 dpf zebrafish nosepits shows decreased numbers of both motile and primary cilia in *togaram1*-/- (-21) as compared to wildtype. **(E)** Wild-type **(F)** and *togaram1*-/- (-21) immunofluorescence of hindbrain ventricles show a clear decrease in cilia number and acetylation in *togaram1*-/- (-21) compared to wildtype. Scale bars for (d – f) represent 10 µm. **(G)** Quantification of acetylated tubulin of cilia in hindbrain ventricles, p-values were determined by student's t-test.

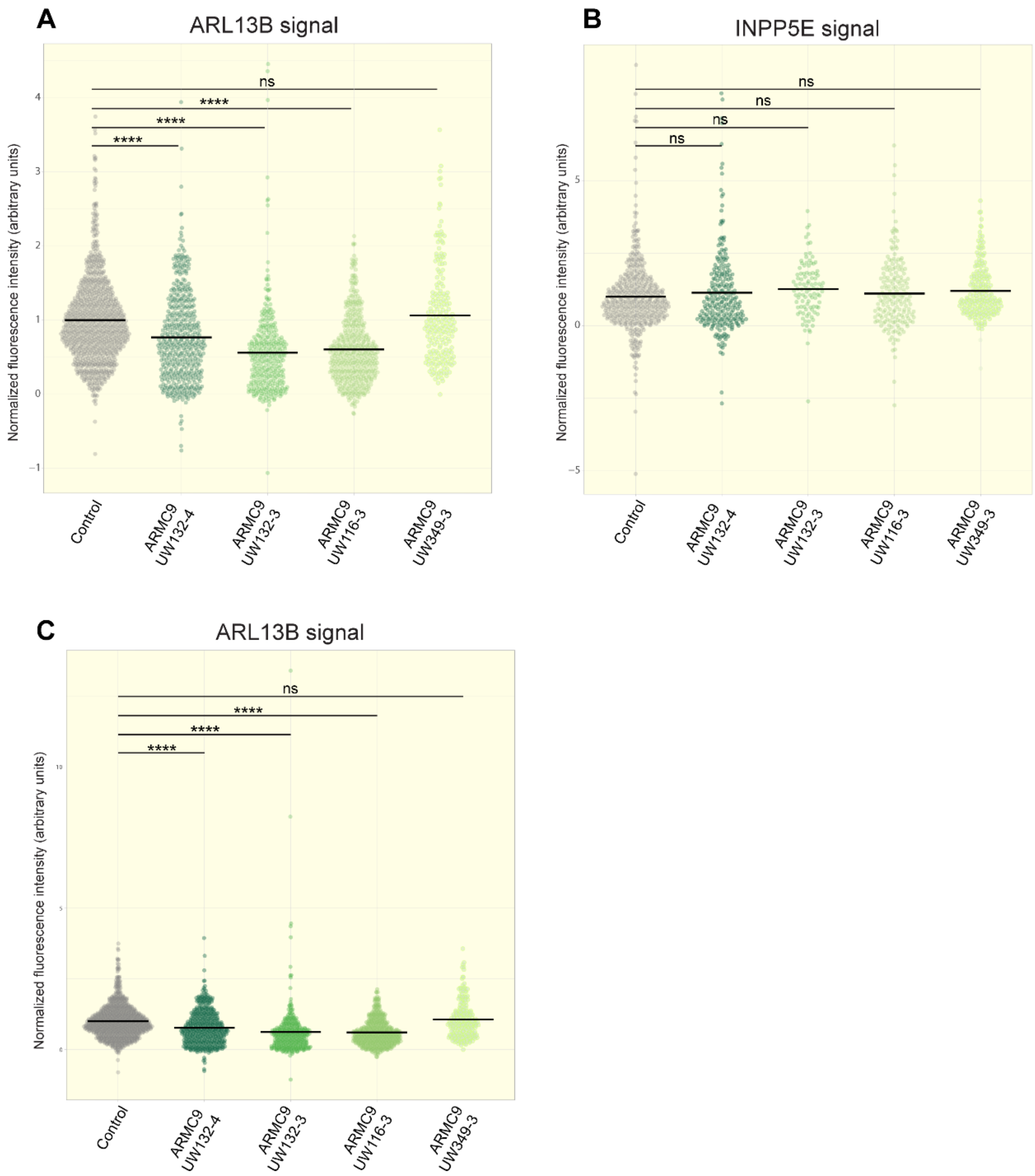


**Supplemental Figure 7** Ciliary INPP5E levels are maintained across *ARMC9* patient fibroblast lines. **(A)** Measurements of normalized fluorescence intensity of ARL13B in arbitrary units along the ciliary length in control and *ARMC9* patient-derived fibroblasts. >200 cilia were assessed per line for measurements. (control = 935 cilia, UW132-4 = 472 cilia, UW132-3 = 337 cilia, UW116-3= 477 cilia, and UW349-3= 208 cilia). p<0.0001 between control and ARMC9 patient cilia UW132-4, UW132-3, UW116-3, and results were not significant for control versus UW349-3 (p=0.5140) by one-way ANOVA. **(B)** Measurements of normalized fluorescence intensity of INPP5E in arbitrary units along the ciliary length in control and ARMC9 patient-derived fibroblasts. >99 cilia were assessed per line for measurements. (control = 471 cilia, UW132-4 = 248 cilia, UW132-3 = 100 cilia, UW116-3= 180 cilia, and UW349-3= 194 cilia). Results were not significant as assessed by one-way ANOVA. Dunett's multiple comparison test yielded the following p values: control versus UW132-4 p= 0.4373, UW132-3 p=0.1701, UW116-3 p=0.7163, UW349-3 p=0.2038. **(C)** In the dot plots shown in (A), data point above 4 and below -2 have been excluded from this graph for clarity, but retained in the statistical analysis, (C) shows the complete graph of all data points. Symbols indicate the following P values: ns, P > 0.05; *, P ≤ 0.05; **, P ≤ 0.01; ***, P ≤ 0.001; ****, P ≤ 0.0001.

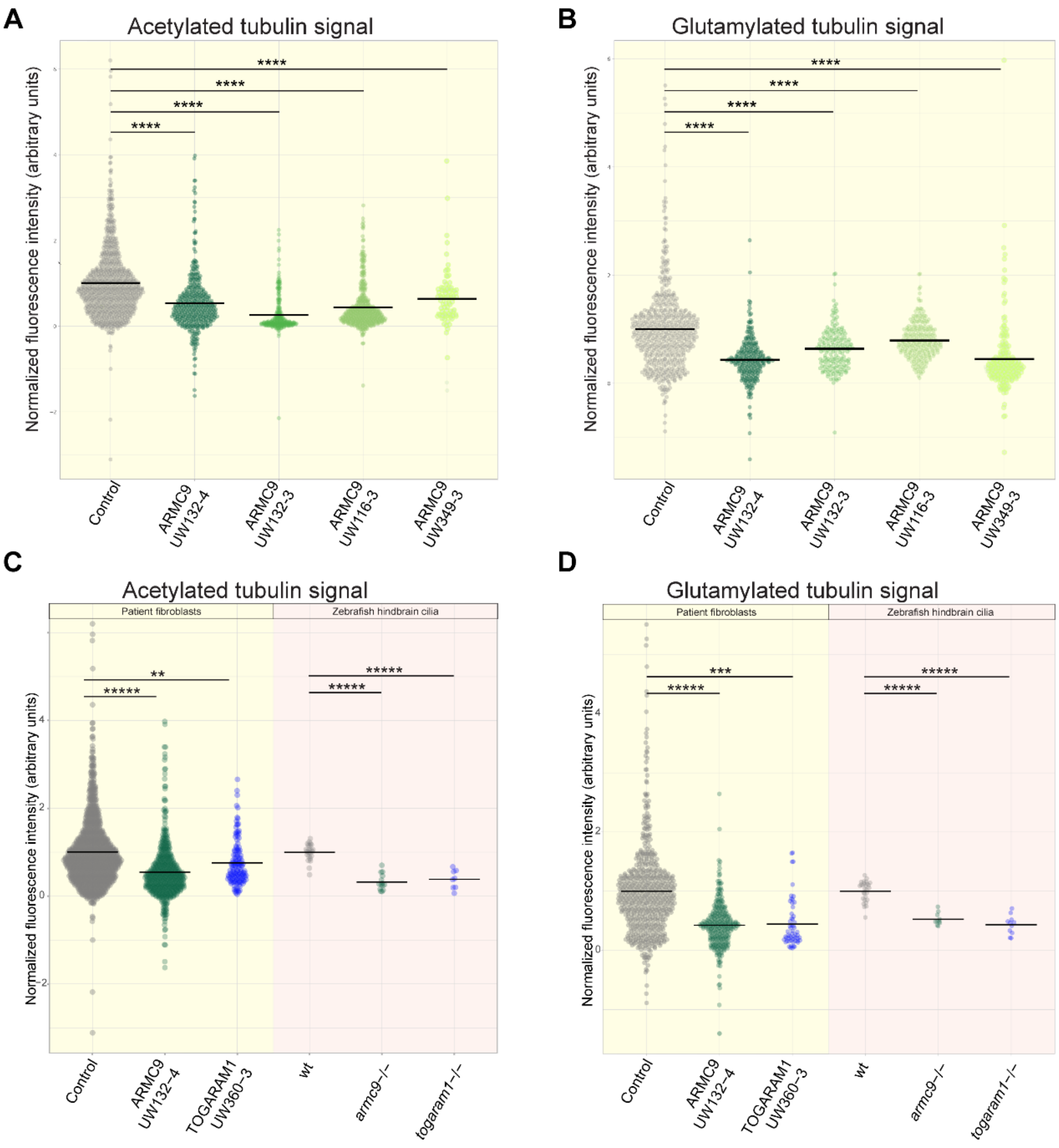


**Supplemental Figure 8** Aberrant post-translational modifications across *ARMC9* patient fibroblast lines. **(A)** Measurements of normalized fluorescence intensity of acetylation in arbitrary units along the ciliary length in control and *ARMC9* patient-derived fibroblasts. >80 cilia were assessed per line for measurements. (control = 848 cilia, UW132-4 = 397 cilia, UW132-3 = 258 cilia, UW116-3= 399 cilia, and UW349-3= 84 cilia). p<0.0001 between control and ARMC9 patient cilia by one-way ANOVA. **(B)** Measurements of normalized fluorescence intensity of polyglutamylation in arbitrary units along the ciliary length in control and *ARMC9* patient-derived fibroblasts. >150 cilia were assessed per line for measurements. (control = 557 cilia, UW132-4 = 262 cilia, UW132-3 = 179 cilia, UW116-3= 253 cilia, and UW349-3= 194 cilia). p<0.0001 between control and ARMC9 patient cilia by one-way ANOVA. **(C)** Full graphs of the dot plots shown in Figure 7 E, **(D)** and F showing all data points. Symbols indicate the following P values: ns, P > 0.05; *, P ≤ 0.05; **, P ≤ 0.01; ***, P ≤ 0.001; ****, P ≤ 0.0001.

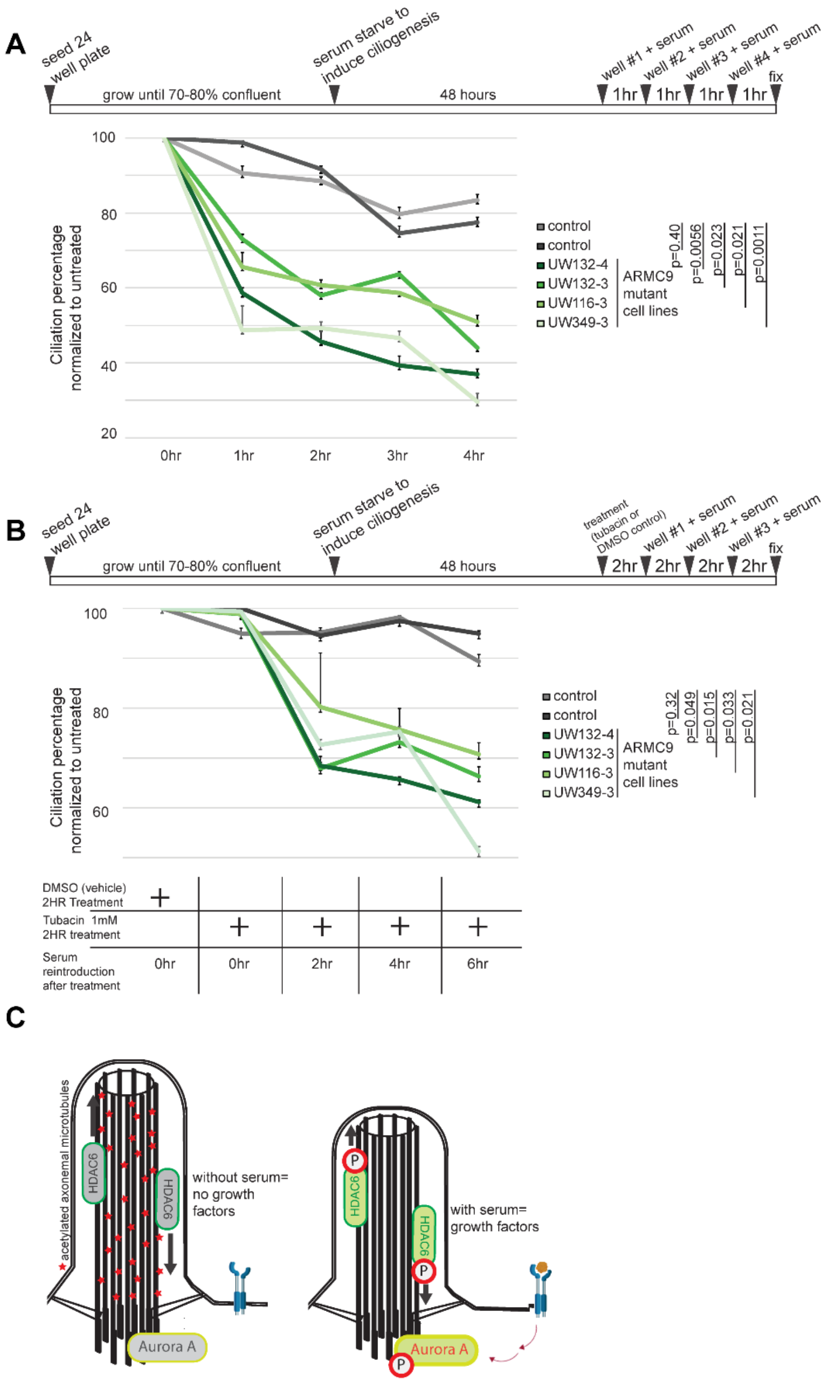


**Supplemental Figure 9** *ARMC9* patient fibroblasts exhibit reduced stability. **(A)** Serum readdition assay schematic and time course showing ciliation percentages normalized to serum-starved cells for each cell line. Note accelerated loss of cilia in all *ARMC9* lines. **(B)** Serum readdition assay schematic with HDAC6 inhibitor (Tubacin) treatment to block HDAC6 activity (to test whether the faster resorption observed in ARMC9 lines observed in A was caused by overactive deacetylating enzyme) and time course showing ciliation percentages normalized to vehicle treated serum-starved cells for each cell line. Error bars represent confidence intervals and p-values were determined by student's t-test with unequal variance for both (A and B). Note HDAC6-inhibition does not rescue accelerated loss of cilia in *ARMC9* lines, but does inhibit resorption in controls. **(C)** Schematic model of HDAC6 activity in ciliary disassembly. Upon initiation of resorption, histone deacetylase 6 (HDAC6) becomes activated and deacetylates the ciliary microtubules, a required step for resorption (67).

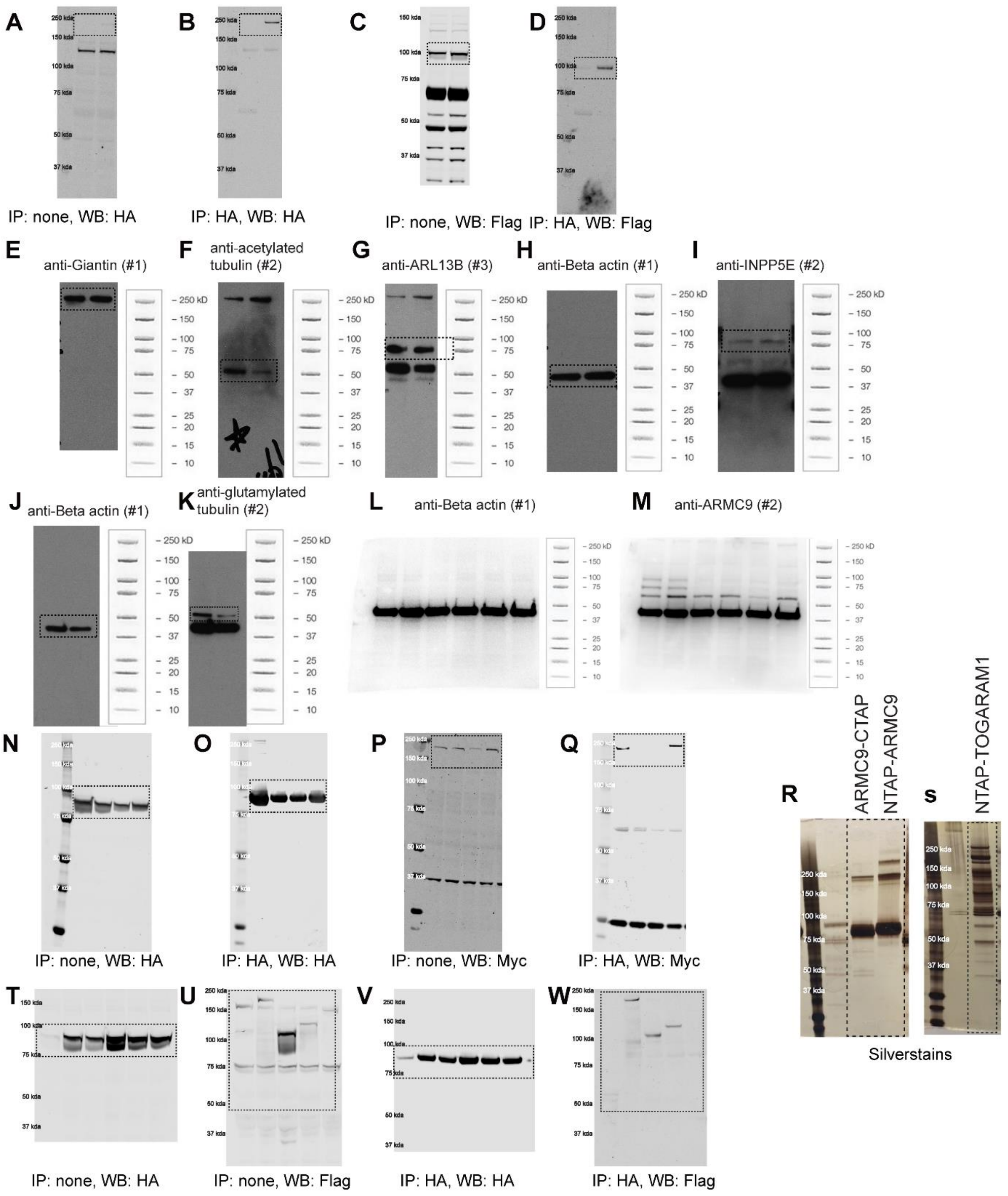


**Supplemental Figure 10** Full scans of western blots and silver-stained gels. Dotted boxes indicate the respective crops in the indicated figures. **(A)** HA staining full blot from figure 1 C. **(B)** HA staining full blot from HA IP in figure 1 C. **(C)** Flag staining full blot from figure 1 C. **(D)** Flag staining full blot post HA IP from figure 1 C. **(E)** Giantin staining full blot from figure 6 C, figure 7 A. **(F)** acetylated tubulin staining full blot from figure 7 A. **(G)** ARL13B staining full blot from figure 6 C. **(H)** Beta actin staining full blot from figure 6 D. **(I)** INPP5E staining full blot from figure 6 D. **(J)** Beta actin staining full blot from figure 7 B. **(K)** Polyglutamylated tubulin staining full blot from figure 7 B. **(L)** Beta actin staining full blot from figure 5 A. **(M)** ARMC9 staining full blot from figure 5 A. **(N)** HA staining full blot from figure 3 G. **(O)** HA staining full blot from figure 3 G. **(P)** Myc staining full blot from figure 3 G. **(Q)** Myc staining full blot from figure 3 G. **(R)** Silver stain from figure 1 B. **(S)** Silver stain from figure 1 B. **(T)** HA staining full blot from Supplemental Figure 1 A. **(U)** Flag staining full blot from Supplemental Figure 1 A. **(V)** HA staining full blot post HA IP from Supplemental Figure 1 A. **(W)** Flag staining full blot post HA IP from Supplemental Figure 1 A.
